# Targeted protein S-nitrosylation of ACE2 as potential treatment to prevent spread of SARS-CoV-2 infection

**DOI:** 10.1101/2022.04.05.487060

**Authors:** Chang-ki Oh, Tomohiro Nakamura, Nathan Beutler, Xu Zhang, Juan Piña-Crespo, Maria Talantova, Swagata Ghatak, Dorit Trudler, Lauren N. Carnevale, Scott R. McKercher, Malina A. Bakowski, Jolene K. Diedrich, Amanda J. Roberts, Ashley K. Woods, Victor Chi, Anil K. Gupta, Mia A. Rosenfeld, Fiona L. Kearns, Lorenzo Casalino, Namir Shaabani, Hejun Liu, Ian A. Wilson, Rommie E. Amaro, Dennis R. Burton, John R. Yates, Cyrus Becker, Thomas F. Rogers, Arnab K. Chatterjee, Stuart A. Lipton

**Author notes:** (S.A.L.).

## Abstract

Prevention of infection and propagation of SARS-CoV-2 is of high priority in the COVID-19 pandemic. Here, we describe S-nitrosylation of multiple proteins involved in SARS-CoV-2 infection, including angiotensin converting enzyme 2 (ACE2), the receptor for viral entry. This reaction prevents binding of ACE2 to the SARS-CoV-2 Spike protein, thereby inhibiting viral entry, infectivity, and cytotoxicity. Aminoadamantane compounds also inhibit coronavirus ion channels formed by envelope (E) protein. Accordingly, we developed dual-mechanism aminoadamantane nitrate compounds that inhibit viral entry and thus spread of infection by S-nitrosylating ACE2 via targeted delivery of the drug after E-protein channel blockade. These non-toxic compounds are active *in vitro* and *in vivo* in the Syrian hamster COVID-19 model, and thus provide a novel avenue for therapy.

---

The process of SARS-CoV-2 infection first involves binding of the viral Spike protein to a cell surface receptor, which has been shown to be ACE2^1^. Viral entry into cells can be accomplished by fusion of the viral envelope (E) protein, located on SARS-CoV-2 near the Spike protein, to facilitate fusion with the cell surface, or endocytosis with subsequent envelope fusion to endosomal membranes^2–4^. We reasoned that using the E protein viroporin channel to target a molecular warhead to the ACE2 receptor to inhibit interaction with the Spike protein could yield a novel mechanism for drug action to treat COVID-19.

Along these lines, we built upon our experience in developing the aminoadamantane drug, memantine, as an FDA-approved treatment for Alzheimer’s disease, and synthesized a number of drugs with improved efficacy^5–9^. These new compounds, termed aminoadamantane nitrates, offer dual-allosteric inhibition of the ion channel associated with the *N*-methyl-D-aspartate (NMDA)-type of glutamate receptor in the brain, with the aminoadamantane moiety providing channel block as well as targeted delivery of a nitric oxide (NO)-related group to S-nitrosylate and thus further inhibit the receptor. Moreover, these aminoadamantane nitrates have displayed no untoward side effects, such as hypotension or other NO-associated actions, in two-species toxicity studies^5–9^. Intriguingly, aminoadamantane drugs like amantadine and memantine were originally developed as anti-viral agents because they also block the ion channel found in the envelope of multiple viruses, including influenza and the β-coronaviruses, and anecdotal reports in humans suggest that they may possibly offer some efficacy for SARS-CoV-2 but definitive data are lacking^2, 3, 10–14^. Coupled with the facts that SARS-CoV and SARS-CoV-2 have also been shown to be susceptible to NO, in part by inhibiting their protease and replication cycle^15, 16^, and NO-based therapies have shown promise in human clinical trials for COVID-19 treatment^17, 18^, we postulated that the new aminoadamantane nitrate drugs might provide mechanistic information on the mode of action of NO against SARS-CoV-2 and offer improved anti-viral activity. Indeed, we show here that the aminoadamantane moiety can block the ion channel in the envelope of SARS-CoV-2 to provide a guided missile to target a therapeutic warhead to ACE2 and thus prevent interaction with the nearby Spike protein. Accordingly, we demonstrate that the viral receptor, ACE2, can be S-nitrosylated by NO-related species generated by the nitro adduct of the aminoadamantane nitrate compounds to inhibit viral entry.

## Results

### S-Nitrosylation of ACE2 inhibits binding of SARS-CoV-2 Spike protein

Initially, we investigated the molecular mechanism whereby NO-related species might inhibit SARS-CoV-2 activity. As assessed by biotin-switch assay, we found that the host cell membrane protein receptor for SARS-CoV-2, ACE2, and a protease that cleaves the viral Spike (S) protein, transmembrane serine protease 2 (TMPRSS2), both of which are necessary for viral entry and infectivity^1, 19^, could be S-nitrosylated by the physiological small molecule NO donor and transnitrosylating agent S-nitrosocysteine (SNOC) (Fig. 1a–c). Interestingly, multiple cysteine residues have been shown to be of importance in ACE2 and TMPRSS2 activity, so S-nitrosylation might be expected to disrupt their activity^20, 21^.

**Fig. 1.**
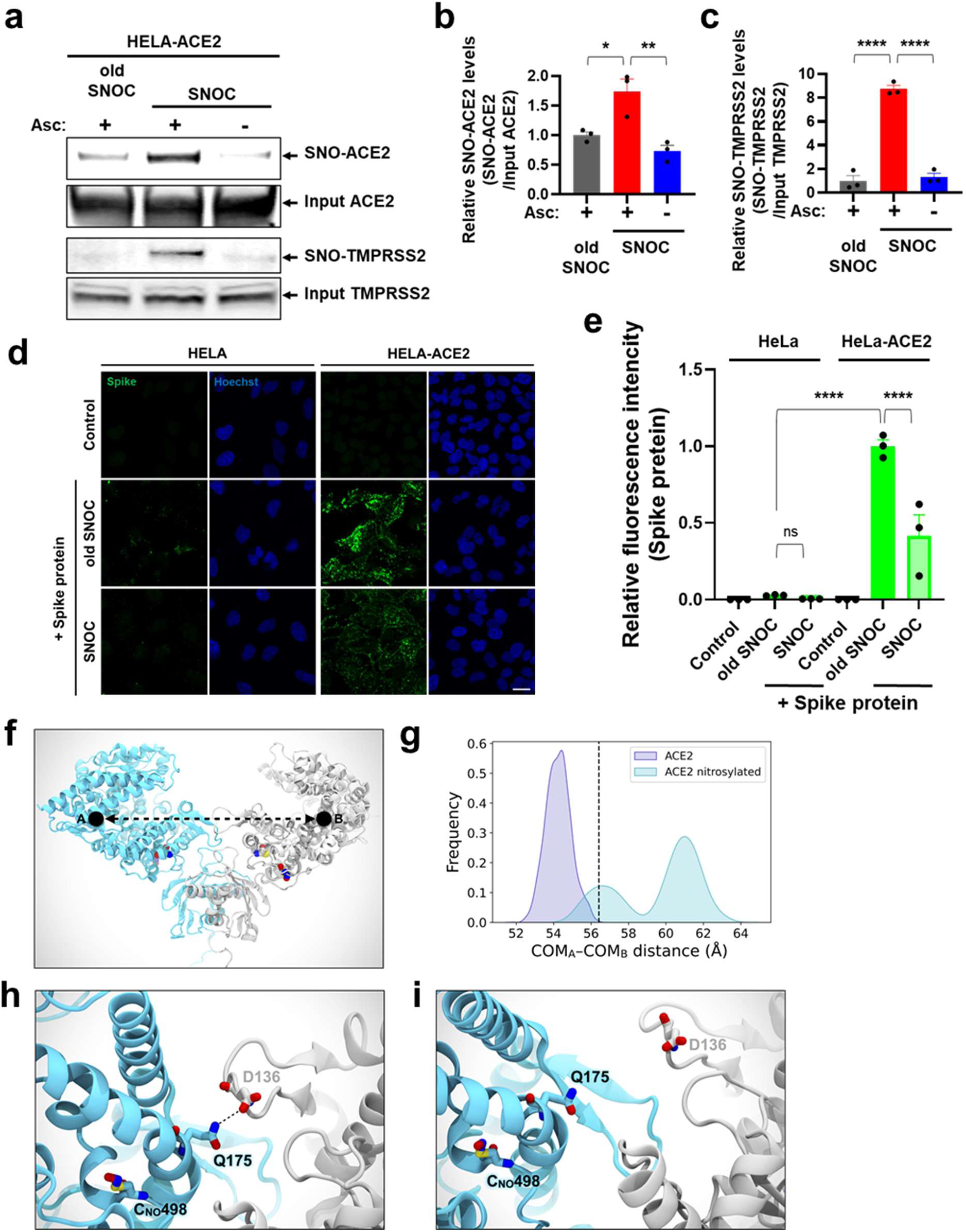
SNOC increases S-nitrosylation of ACE2 and inhibits binding of SARS-CoV-2 Spike (S) protein. **a**, Assay for SNO-ACE2 and SNO-TMPRSS2 in HeLa-ACE2 cells. Cells were exposed to 100 μM SNOC or, as a control, ‘old’ SNOC (from which NO had been dissipated). After 20 minutes, cell lysates were subjected to biotin-switch assay to assess S-nitrosylated (SNO-) and input (total) proteins detected by immunoblotting with cognate antibody. The ascorbate minus (Asc-) sample served as a negative control. **b**, **c**, Ratio of SNO-ACE2/input ACE2 protein and SNO-TMPRSS2/input TMPRSS2 protein. Data are mean + s.e.m., \**P* < 0.05, \*\**P* < 0.01, \*\*\**P* < 0.001 by ANOVA with Tukey’s multiple comparisons. *n* = 3 biological replicates. **d**, HeLa and HeLa-ACE2 cells were pre-exposed to 100 μM SNOC or old SNOC. After 30 minutes, 10 μg/ml of purified recombinant SARS-CoV-2 Spike (S1+S2) protein was incubated with the cells. After 1 h, cells were fixed with 4% PFA for 15 minutes, and bound Spike protein was detected by anti-Spike protein antibody; nuclei stained with 1 μg/ml Hoechst. Cells were imaged by confocal fluorescence microscopy. Scale bar, 20 μm. **e**, Quantification of relative fluorescence intensity. Data are mean + s.e.m., \*\*\*\**P* < 0.0001 by ANOVA with Tukey’s multiple comparisons. *n* = 3 biological replicates**. f**, Molecular representation of the S-nitrosylated-ACE2/RBD model upon transient detachment at the level of the peptidase domain dimeric interface. SNO-Cys261 and SNO-Cys498 are shown with Van der Waals spheres. The black dots indicate qualitative placement of centers of mass (COM) for each ACE2 protomer, and the dashed arrow represents the distance between COMs. Spike’s RBDs and N-glycans, which were included in the simulation, are hidden for image clarity. SpBD, Spike binding domain; CLD, collectrin-like domain; PD, peptidase domain. **g**, Distribution of the distance between COMs from molecular dynamics simulations of WT ACE2/RBD (purple) vs. nitrosylated-ACE2/RBD (cyan). Dashed black line at approximately 56.5 Å indicates the reference distance between COMs calculated from the cryo-EM structure (PDB: 6M17). S-Nitrosylated-ACE2/RBD shows an overall larger distance between COMs with a bimodal distribution. **h**, Close-up image illustrating Q175A to D136B interaction present in starting conformations of the S-nitrosylated-ACE2 system. **i**, Close-up image illustrating the disruption of the interaction between Q175A and D136B occurring along the dynamics of the S-nitrosylated-ACE2 system.

We focused on S-nitrosylation of ACE2 (forming SNO-ACE2), reasoning that this nitrosylation reaction might prevent binding of SARS-CoV-2 S protein to ACE2, thus inhibiting viral infection. To test this premise, we exposed HeLa cells stably expressing human ACE2 (HeLa-ACE2) to SNOC and assessed SNO-ACE2 formation by biotin-switch assay. To evaluate binding of the S protein to these HeLa-ACE2 cells, we then incubated the cells with purified recombinant SARS-CoV-2 Spike protein (S1+S2). Since NO dissipates very quickly from SNOC (<5 minutes at neutral pH), and Spike protein was added sequentially after this period, we could rule out the possibility of direct S-nitrosylation of Spike protein by SNOC under these conditions^22, 23^. We found that the formation of SNO-ACE2 was stable for at least 12 h (Extended Data Fig. 1). The receptor binding domain (RBD) in the S1 subunit of the SARS-CoV-2 Spike glycoprotein binds to ACE2 expressed on the surface of host cells, while the C-terminal S2 membrane anchoring subunit functions to translocate virus into host cells^21, 24^. After preincubation of HeLa-ACE2 cells with SNOC, we found significantly decreased binding of purified S protein to HeLa-ACE2-cells (Fig. 1d, e), consistent with the notion that the cysteine residue(s) susceptible to S-nitrosylation in ACE2 affected S protein binding.

Human ACE2 protein contains eight cysteine residues, six of which participate in formation of three pairs of disulfide bonds, and the remaining two (Cys^2^^61^ and Cys^498^) are present as free thiols (or thiolates) (Extended Data Fig. 2a)^21^ and thus potentially available for S-nitrosylation via reversible nucleophilic attack on a nitroso nitrogen to form an SNO-protein adduct^25^. Accordingly, we performed site-directed mutagenesis of these cysteine residues in ACE2 and found that C261A, C498A, or C261A/C498A mutation significantly inhibited SNOC-mediated S-nitrosylation on biotin-switch assays, consistent with the notion that these two cysteine residues are targets of S-nitrosylation (Extended Data Fig. 2b, c). Moreover, mass spectrometry confirmed the presence of S-nitrosylated ACE2 at Cys^261^ and Cys^498^ after exposure to SNOC (Extended Data Fig. 2d).

**Fig. 2.**
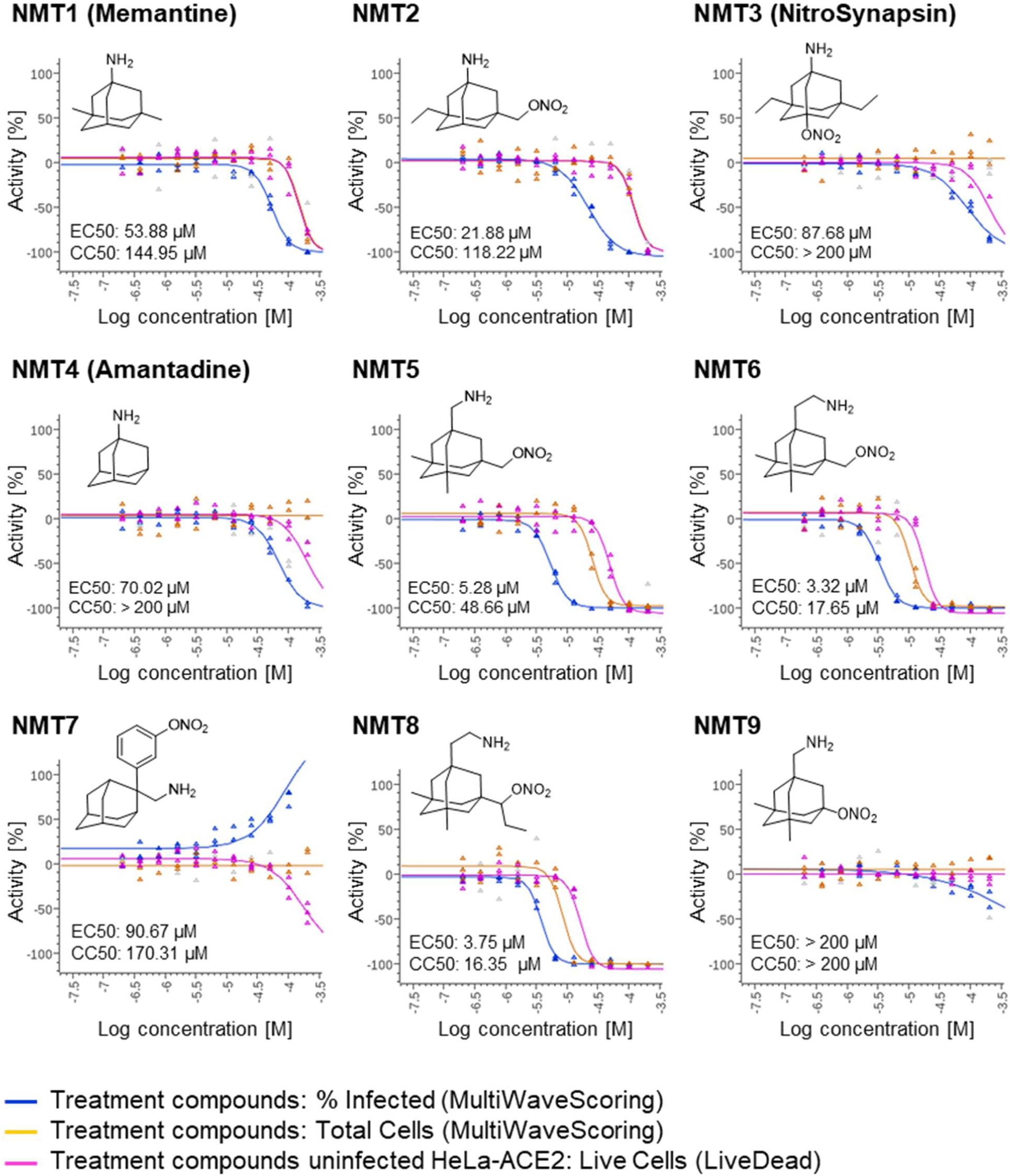
Dose-response of drugs screened against SARS-CoV-2. Dose-response curves showing the EC_50_ of each compound against SARS-CoV-2 (% infected cells, blue), total cell counts (orange) in the infection experiment and the CC_50_ for uninfected host cell toxicity (magenta), as assessed in HeLa-ACE2 cells. See also Extended Data Table 1 for full dataset.

### S-Nitrosylation of ACE2 destabilizes dimer formation and thus Spike protein binding

Notably, these S-nitrosylation sites are located near the collectrin-like domain (CLD) region rather than the Spike protein-binding domain region of ACE2 (Fig. 1f). This suggests that S-nitrosylation may affect the conformation of ACE2 protein at some distance from the S-nitrosylated cysteine residue(s) to diminish binding of ACE2 to trimeric S protein^21, 24, 26^. Accordingly, explicitly solvated, all-atom molecular dynamics simulations of the S-nitrosylated-ACE2/RBD complex in plasma membrane show that the distance between each S-nitrosylated ACE2 protomer’s center of mass is overall much longer and more broadly distributed than in simulations of wild-type (WT) ACE2 dimer (Fig. 1f, g)^26^. This behavior indicates a certain extent of destabilization of the dimer interface imparted by S-nitrosylation, particularly of C^498^. Specifically, at the beginning of the simulations, the S-nitrosylated-ACE2/RBD model displays a hydrogen bond between Q175_A_ and Q139_B_, which is then interchanged with D136_B_ (Fig. 1h). This is the only interaction between the peptidase domains (PD) of the two protomers, as also reported for the initial cryo-EM structure^27^. Importantly, over the course of our simulations, this interaction was progressively lost (Fig. 1i), leading to partial disruption of the PD dimeric interface and transient detachment of the two protomers. Therefore, we hypothesize that the addition of S-nitrosylation at the side chain of C^498^, which is located in the vicinity of Q^175^, could be sufficient to induce rearrangement in the packing of secondary structural elements of this region, leading in turn to the disruption of the only point of contact between the two PDs of ACE2. The loss of this contact may potentially trigger a further destabilization at the level of the dimeric interface between the neck domains. Alteration of ACE2 dimer stability has the potential to interfere with the SARS-CoV-2 Spike binding^28^, thus abrogating infection.

### Screening aminoadamantane nitrate compounds against SARS-CoV-2 infection

Next, we examined the effect of aminoadamantane nitrate compounds on SARS-CoV-2 entry into cells, causing infection. Aminoadamantanes have been reported to directly bind to the viroporin ion channel formed by the SARS-CoV-2 envelope (E) protein^10, 29, 30^. Therefore, we screened our series of aminoadamantane and nitro-aminoadamantane compounds^6–8^ as potential therapeutic drugs against SARS-CoV-2 — these latter drugs might be expected to bind to the viral channel, thus targeting S-nitrosylation to ACE2 to inhibit its interaction with Spike protein and thus viral entry. Specifically, we tested in a masked fashion the efficacy against live SARS-CoV-2 in HeLa-ACE2 cells of aminoadamantanes (memantine, blindly coded as NMT1, and amantadine/NMT4) and aminoadamantane nitrate compounds (NMT2, NMT3 and NMT5-NMT9) (Fig. 2, full data set shown in Extended Data Table 1). As positive controls, we used remdesivir, apilimod, and puromycin (Extended Data Table 1)^31, 32^. In determining the therapeutic potential of these compounds, we considered the selectivity index (SI) that compares a compound’s half-maximal non-specific cytotoxicity (CC_50_) in the absence of infection to its half-maximal effective antiviral concentration (EC_50_) (CC_50_/EC_50_) (Extended Data Table 1). The SI can be considered an *in vitro* indicator of therapeutic index and ideally would approach 10. The aminoadamantane compounds alone (amantadine and memantine) offered no efficacy, and thus were not studied further. In contrast, several of the aminoadamantane nitrate compounds offered some degree of protection from infection. However, NMT6 and NMT8 may have done this simply by killing the host cells irrespective of infection, as evidenced by its off-target killing of uninfected cells in the live/dead assay (Fig. 2, Extended Data Table 1). Among the 7 aminoadamantane nitrate compounds tested, NMT5 displayed the best combination of EC_50_ and CC_50_ (SI = 9.2) with an EC_50_ for protection against SARS-CoV-2 of 5.28 μM (Fig. 2, Extended Data Table 1); this concentration of compound is well within the micromolar amounts attainable in human tissues at well-tolerated doses, as tested in two animal species^6–9, 33^. Additionally, NMT3 (also known as NitroSynapsin), which was already being developed for CNS indications^6–9^ displayed some degree of protection against SARS-CoV-2 with an EC_50_ of 87.7 μM, although this value may be artificially high due to the short half-life of NMT3 in aqueous solution under *in vitro* conditions^6, 9, 34^. Hence, these two compounds were advanced for further study. We next asked if NMT3 and NMT5 could S-nitrosylate ACE2. We found that NMT5 > NMT3 effectively S-nitrosylated ACE2 both *in vitro* in HeLa-ACE2 cells and *in vivo* in Syrian hamsters, as assessed by the biotin-switch assay (Fig. 3a, b, Extended Data Fig. 3). Notably, a statistically significant increase in the level of S-nitrosylated ACE2 was observed in the SARS-CoV-2 target tissues of lung and kidney at 48 h after oral administration of a single dose of drug at 10 mg/kg (Extended Data Fig. 3d–i). Consistent with the structure-activity relationship (SAR) indicating that SNO-ACE2 was associated with the anti-viral effect of NMT5 and NMT3, the other aminoadamantane nitrates (including NMT6 and NMT8) did not S-nitrosylate ACE2 at low micromolar concentrations (Extended Data Fig. 4).

**Fig. 3.**
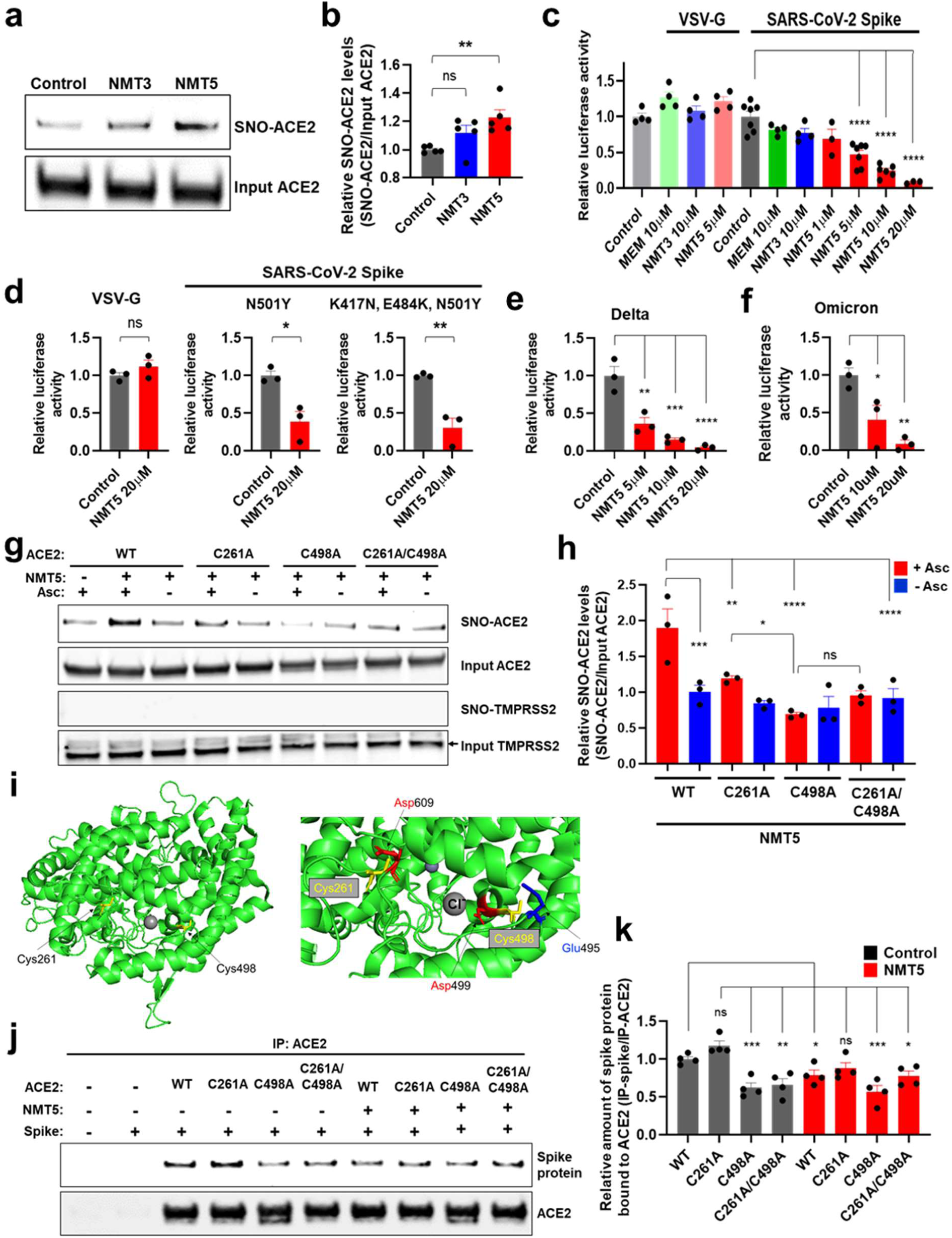
NMT5 S-nitrosylates ACE2 and inhibits SARS-CoV-2 pseudoviral entry. **a**, HeLa-ACE2 cells were treated with 10 μM NMT3 or 5 μM NMT5. After 1 h, cell lysates were subjected to biotin-switch assay for protein S-nitrosylation, detected by immunoblotting with anti-ACE2 antibody. **b**, Ratio of SNO-ACE2/input ACE2 protein. Data are mean + s.e.m., \*\**P* < 0.01, ns: not significant by ANOVA with Tukey’s multiple comparisons. *n* = 4 biological replicates. **c**, HeLa-ACE2 cells were incubated with SARS-CoV-2 Spike (D614) or VSV-G (control) pseudovirus particles in the presence and absence of MEM (memantine), NMT3, or NMT5. After 48 h, viral transduction efficiency was monitored by luciferase activity. Data are mean + s.e.m., \*\*\*\**P* < 0.0001 by ANOVA with Tukey’s multiple comparisons. *n* = 3 to 6 biological replicates. **d**–**f**, HeLa-ACE2 cells were incubated in the presence and absence of NMT5 with SARS-CoV-2 N501Y Spike, SARS-CoV-2 K417N/E484K/N501Y Spike, or VSV-G (control) pseudovirus particles (d); or with SARS-CoV-2 delta variant (e) or omicron variant pseudovirus particles (f). After 48 h, viral transduction efficiency was monitored by luciferase activity. Data are mean + s.e.m., \**P* < 0.05, \*\**P* < 0.01, \*\*\**P* < 0.001, \*\*\*\**P* < 0.0001 by two-tailed Student’s *t* test for single comparison (d) or ANOVA with Tukey’s for multiple comparisons (e, f). ns: not significant, *n* = 3 biological replicates. **g**, HEK293T cells were transfected with plasmids encoding human WT ACE2 or non-nitrosylatable mutant ACE2 (C262A, C498A, or C261A/C498A). One day later, cells were treated with 10 μM NMT5, and 1-h later subjected to biotin-switch assay for detection of S-nitrosylated proteins by immunoblotting with anti-ACE2 and anti-TMPRSS2 antibodies. The absence of ascorbate (Asc-) served as a negative control. **h**, Ratio of SNO-ACE2/input ACE2. Data are mean + s.e.m., \**P* < 0.05, \*\**P* < 0.01, \*\*\**P* < 0.001, \*\*\*\**P* < 0.0001 by ANOVA with Fisher’s LSD multiple comparisons. ns: not significant, *n* = 3 biological replicates. **i**, Crystal structure of ACE2 (left panel; PDB ID: 6M0J) with enlarged view of S-nitrosylation sites of ACE2. Glu^495^ and Asp^499^, acidic amino-acid residues, surround Cys^498^ (right panel). **j**, HEK293T cells were transfected with plasmids encoding human WT ACE2 or non-nitrosylatable mutant ACE2 (C262A, C498A, or C261A/C498A). One day later, cells were exposed to 1 μg/ml of purified recombinant SARS-CoV-2 Spike protein in the presence or absence of 5 μM NMT5; after 1 h, cells were lysed and subjected to co-IP with anti-ACE2 antibody. Immunoprecipitated ACE2 and Spike protein were detected by immunoblotting with anti-ACE2 and anti-Spike protein antibodies. **k**, Ratio of IP-ACE2/IP-Spike protein. Data are mean + s.e.m., \**P* < 0.05, \*\**P* < 0.01, \*\*\**P* < 0.001, ns: not significant by ANOVA with Fisher’s LSD multiple comparisons. *n* = 5 biological replicates.

**Fig. 4.**
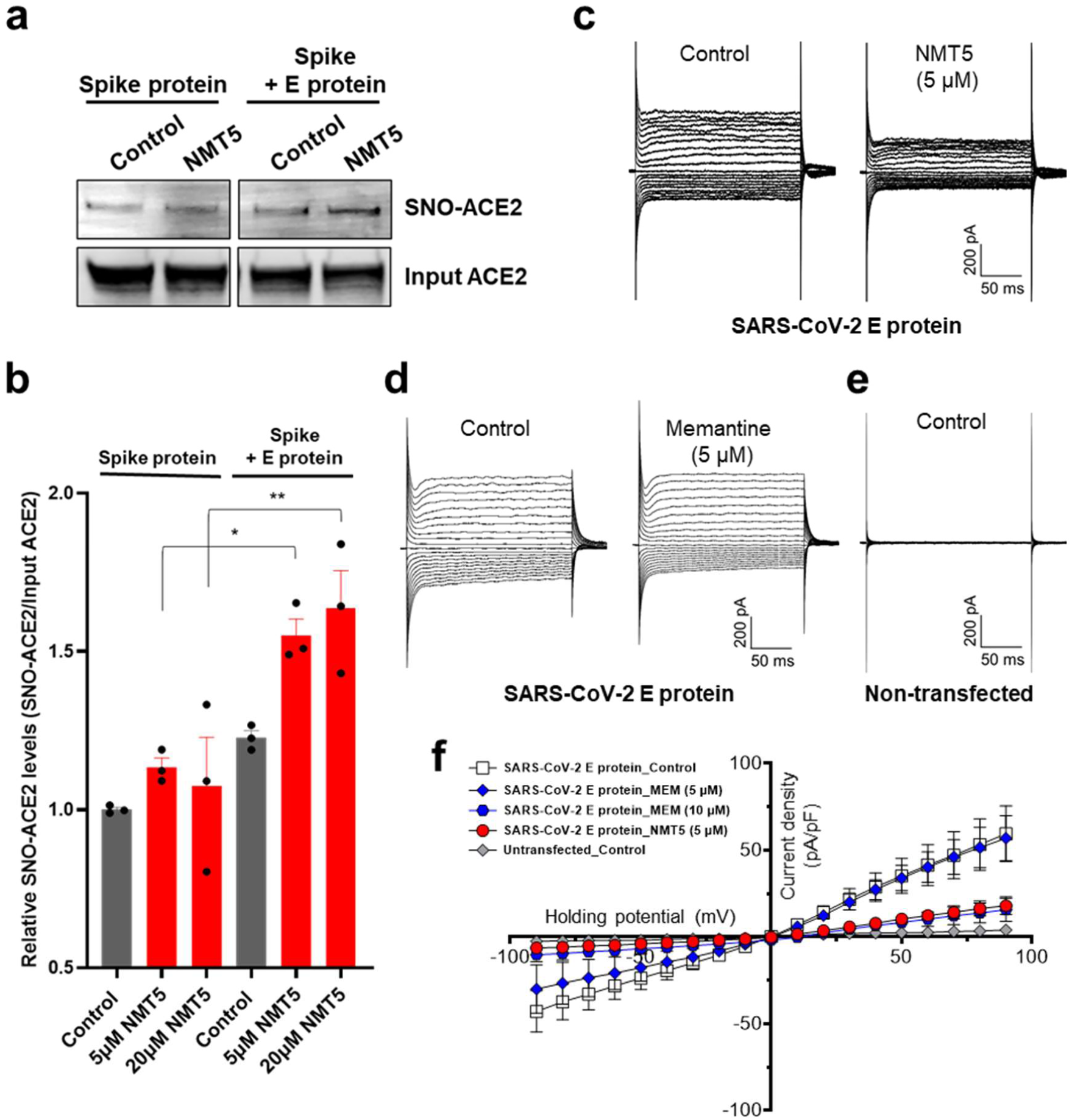
Targeted S-nitrosylation of ACE2 and Inhibition of envelope (E) viroporin protein channel by NMT5. **a**, E protein plasmid was transiently transfected into HEK293-Spike protein cells. After 1 day, cells were harvested and plated onto HeLa-ACE2 cells in the presence or absence of 5 µM NMT5. After 30 min, cell lysates were subjected to biotin-switch assay to monitor protein S-nitrosylation of ACE2, detected by immunoblotting. **b**, Ratio of SNO-ACE2/total input ACE2 protein. Data are mean + s.e.m., \**P* < 0.05, \*\**P* < 0.01 by ANOVA with Tukey’s multiple comparisons. *n* = 3 biological replicates. **c**–**e**, Representative traces of whole-cell currents from untransfected (*n* = 4), and transiently transfected (*n* = 15) HEK293T cells before and after application of memantine or NMT5 during patch-clamp recording. Whole-cell currents were generated by holding cells at 0 mV and applying voltage steps between −90 and +90 mV in increments of 10 mV. **f**, Current-voltage (*I*–V) curves from steady-state current density (pA/pF) versus holding potential (mV) for memantine (MEM, 5 and 10 µM) and NMT5 (5 µM).

### Drug candidate NMT5 can S-nitrosylate ACE2 and inhibit entry of SARS-CoV-2 variants in pseudovirus assays

Since we had found that S-nitrosylation of ACE2 inhibited the binding of SARS-CoV-2 Spike protein, we next asked if NMT3- or NMT5- mediated SNO-ACE2 formation could prevent viral entry into host cells. To test this premise, we employed a replication-deficient Maloney murine leukemia virus (MLV)- based SARS-CoV-2 Spike protein pseudotype virus, initially using the most prevalent strain of Spike protein (D614) as of early 2020^35^. We examined whether NMT3 and NMT5 could suppress infection with this SARS-CoV-2 pseudovirus. We found that NMT5 inhibited SARS-CoV-2 pseudoviral entry in a dose-dependent manner, with 5 µM inhibiting 53%, 10 µM 76%, and 20 µM 92% (Fig. 3c). NMT3 showed more limited ability to suppress pseudovirus entry, ∼24% at 10 µM. The fact that S-nitrosylation of ACE2 manifested inhibition in the pseudovirus assay (as shown in Fig. 3c) at the approximately the same EC50 of 5 µM as found in the live virus infection assay (Fig. 2) strongly implies that SNO-ACE2 formation is indeed the predominant mechanism by which NMT5 prevents viral infection. As a control, the NMT5 metabolite lacking the nitro group did not suppress SARS-CoV-2 infection in the pseudovirus entry assay (Fig. 3c, Extended Data Fig. 5). As a further control, the pseudovirus entry assay performed with vesicular stomatitis virus G protein (VSV-G) was unaffected by NMT3 or NMT5 (Fig. 3c).

**Fig. 5.**
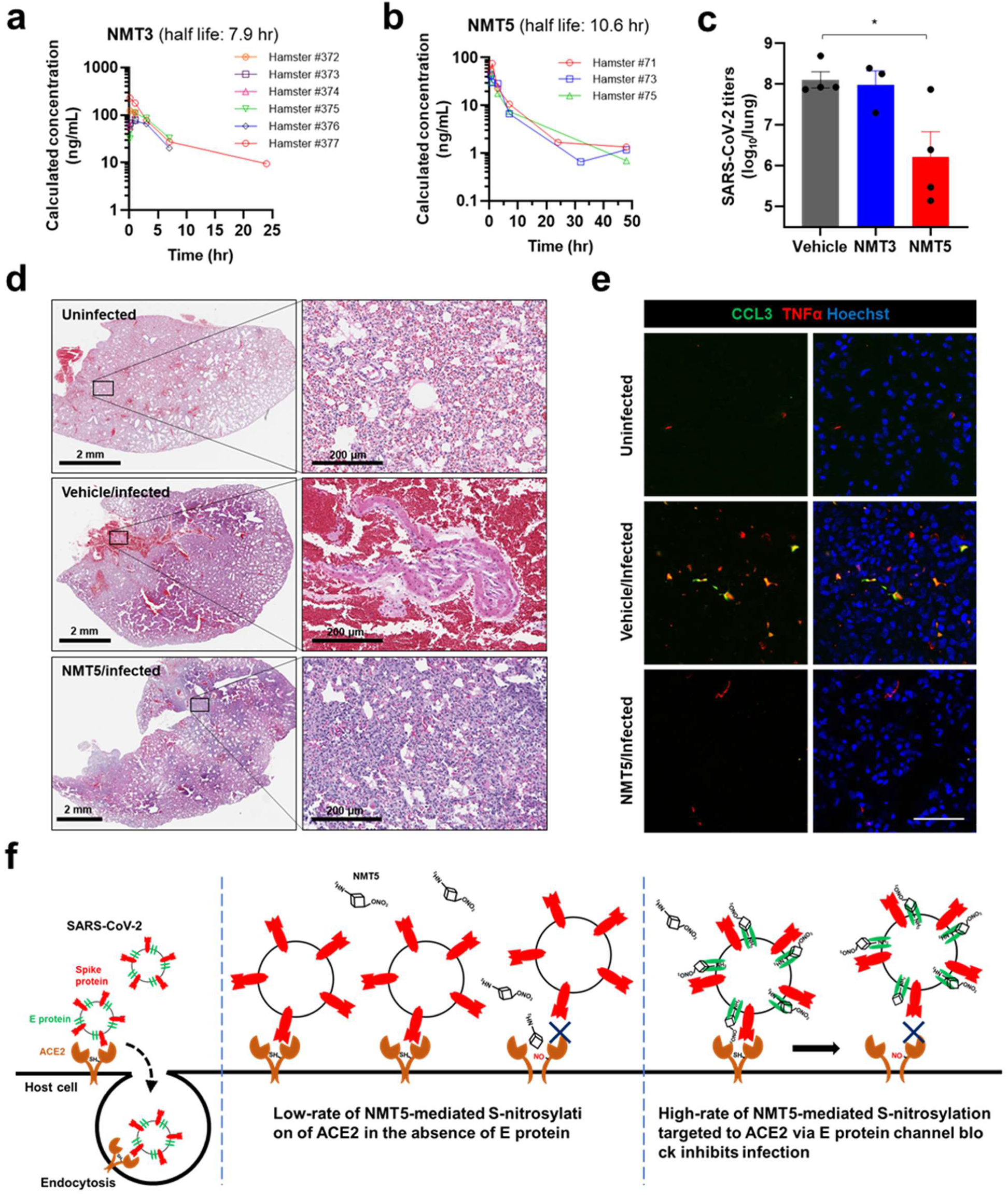
NMT5 inhibits SARS-CoV-2 infection *in vivo* in Syrian hamsters. **a**, **b**, PK data in plasma for NMT3 and NMT5 in Syrian hamsters after an oral dose of 10 mg/kg. **c**, Live viral load in Syrian hamsters monitored by plaque assay from lung tissue 2-d postinfection after treatment with NMT3, NMT5, or vehicle (Control). Data are mean + s.e.m., \**P* = 0.0261 by two-tailed Student’s *t* test (*n* = 8 Syrian hamsters tested). **d**, **e**, Syrian hamsters were sacrificed 5 d after infection with SARS-CoV-2 that were either untreated (labeled Vehicle/Infected) or treated with oral NMT5 (NMT5/Infected), and compared to control (Uninfected). Representative hematoxylin and eosin (H&E)-stained sections showed virtually no areas of large hemorrhage (confluent bright red regions) in NMT5-treated hamster lungs compared to vehicle-treated. Scale bars, 2 mm, high magnification 200 µm (**d**). Representative immunohistochemistry of lung sections stained with anti-TNFα cytokine antibody and anti-CCL3 (MIP-1α) chemokine antibody from untreated- and NMT5-treated hamsters 5-d post SARS-CoV-2 infection; uninfected sections are shown as a control. Merged image with Hoechst stain for DNA shown in *right-hand* panels. Scale bar, 40 µm (**e**). **f**, Schematic of NMT5 targeting of SNO-ACE2 via SARS-CoV-2 E protein. S-Nitrosylation of ACE2 on the host cell inhibits SARS-CoV-2 entry and thus infection.

We next examined if NMT5 could suppress viral infection from more recently identified SARS-CoV-2 variants, including N501Y Spike protein, a common mutation in the B.1.1.7 (or alpha variant, United Kingdom), and B.1.351 (or beta variant, South Africa), P.1 (or gamma variant, Brazil) encountered in the winter of 2020/2021. Additionally, we tested K417N, E484K, N501Y Spike protein, as found in the B.1.351 and P.1 variants, the B.1.617.2 variant (or delta, Indian; T19R, G142D, 156/157 DELETION, R158G, L452R, T478K, D614G, P681R, and D950N), and the BA.1 variant (or omicron, South Africa) found in 2021/2022, which are associated with higher transmissibility or severity as well as altered antigenicity^36^. We found that NMT5 was also effective in reducing infectivity of these SARS-CoV-2 variants, including the delta and omicron variants, by up to 95% (Fig. 3d–f). These results are consistent with the notion that NMT5 >> NMT3-mediated S-nitrosylation of ACE2 can inhibit SARS-CoV-2 entry into host cells.

### NMT5 predominantly S-nitrosylates ACE2 on cysteine residue 498 and inhibits Spike protein binding to ACE2

Next, we sought to determine if NMT5 could modify ACE2 at both of the cysteine residues (Cys^261^ and Cys^498^) that we demonstrated to be susceptible to S-nitrosylation by SNOC. Analysis by cysteine mutation revealed that NMT5 preferentially S-nitrosylated Cys^498^ over Cys^261^ (Fig. 3g, h). Interestingly, the crystal structure of ACE2 shows that an acid/base motif (comprised of Glu^495^ and Asp^499^), which under some conditions may facilitate S-nitrosylation, is present near Cys^498^, while only a partial motif (represented by Asp^609^) is found near Cys^261^ (Fig. 3i)^37–, 39^. This observation is consistent with prior findings that potent or supraphysiological amounts of NO donors such as SNOC can S-nitrosylate cysteine residues surrounded by no motif or only a partial SNO motif^34^, whereas a full SNO motif can facilitate S-nitrosylation by less potent donors, presumably like NMT5. Mechanistically, cysteine thiol groups (in fact, thiolates) are nucleophiles that can perform reversible nucleophilic attack on an electrophilic nitroso nitrogen^40, 41^. The local environment of the thiolate anion can kinetically favor one electrophile over another; moreover, the bulky R-group of NMT5, as an RNOx donor (x = 1 or 2) compared to the small molecule SNOC, could sterically hinder reactivity^34^. Notably, concentrations of NMT5 that significantly inhibited viral entry (∼10 µM) failed to S-nitrosylate other proteins, including TMPRSS2, Spike protein, or E protein (Fig. 3g, Extended Data Fig. 6), demonstrating relative selectivity of NMT5 for ACE2 at the cell surface. While we cannot rule out the possibility that proteins associated with virus intracellular trafficking are S-nitrosylated, this would be less likely as aminoadamantane nitrate compounds are known to act on extracellular rather than intracellular targets^5–9^.

To further investigate the effect of NMT5 on SARS-CoV-2 Spike protein binding to ACE2, we performed co-immunoprecipitation (co-IP) experiments of these two proteins in the presence and absence of NMT5 using anti-ACE2 antibody for IP. As expected, the two proteins co-IP’d, as evidenced on immunoblots. NMT5 (5 µM) significantly diminished this co-IP, consistent with the notion that the drug inhibited the binding of Spike protein to ACE2 (Fig. 3j, k). As controls, the Spike protein was not co-IP’d with cysteine mutant ACE2(C498A) or with double mutant ACE2(C261A/C498A), although mutant ACE2(C261A) was still co-IP’d. These data suggest that S-nitrosylation predominantly of C^498^ of ACE2 is important for Spike protein binding to ACE2. Moreover, NMT5 inhibited co-IP of the Spike protein and ACE2(C261A), while having no effect on mutant ACE2(C498A) or ACE2(C261A/C498A) binding (Fig. 3j, k). Taken together, these results are consistent with the notion that NMT5 inhibits SARS-CoV-2 Spike protein from binding to ACE2 and thus virus entry into the cell via S-nitrosylation of ACE2.

### NMT5 targets S-nitrosylation to ACE2 via blockade of the E-protein viroporin channel

Intriguingly, we found that the presence of the envelope (E) protein of SARS-CoV-2 served to target S-nitrosylation by NMT5 to nearby ACE2 receptor proteins (Fig. 4a, b). To investigate this action further, we assessed the ability of the aminoadamantane compound, memantine, and the lead aminoadamantane nitrate candidate, NMT5, to block ion channel activity of the E protein^11^ using the patch-clamp technique. To test direct interaction with the viroporin channel, we transiently transfected HEK293T cells with a construct encoding the E protein and assessed voltage-dependent currents (vs. uninfected cells) in the presence and absence of drug (Fig. 4c–f). Under our conditions, we found that the presence of the E protein resulted in a robust voltage-dependent current carried by K^+^ that was inhibited in by memantine and with greater potency by NMT5. Notably, the low micromolar concentrations needed to see these effects are within attainable levels in mammalian plasma and tissues, as shown in pharmacokinetic (PK) studies, and have proven to be safe in animal toxicity studies^6-9, 33^.

### NMT5 protects from SARS-CoV-2 infection in the Syrian hamster model of COVID-19

In preparation for *in vivo* drug candidate efficacy testing in a COVID-19 small animal model, we next performed 48-h PK studies after a single oral dose of NMT3 or NMT5 at 10 mg/kg in ∼150 gm Syrian hamsters. We found a half-life in plasma for NMT3 of 7.9 h and for NMT5 of 10.6 h (Fig. 5a, b, Extended Data Table 2). The mean C_max_ for NMT5 was 0.2 µM and ∼0.4 µM for NMT3; NMT3 also displayed a hydroxylated metabolite (full detailed PK dataset shown in Extended Data Tables 3, 4). The fact that NMT5 was found to be more stable than NMT3 by mass spectrometry analysis (Fig. 5a, b, Extended Data Table 2) was also consistent with prior findings^6–9, 33^. Moreover, these drugs are concentrated in tissues up to ∼30-fold over plasma levels. Utilizing a Bayesian-like adaptive clinical trial design, we determined the maximal tolerated dose (MTD) of NMT3 and NMT5 *in vivo* based on dose-ranging toxicity and efficacy studies in 52 Syrian hamsters. To assess treatment efficacy in the Syrian hamster model of COVID-19 at the MTD for NMT5, we administered by oral gavage 200 mg/kg in two equally divided doses separated by 12 h, with the initial dose timed right after challenge with the virus and the second dose 12 h later^35, 42^. Based on the PK results, at this dose, drug levels in tissue should approach or exceed the EC_50_ found in our *in vitro* screens to significantly decrease viral infectivity. We found in the Syrian hamster model that this regimen of NMT5, but not NMT3, knocked down live viral titers of SARS-CoV-2 postinfection by ∼100 fold, as measured by plaque assay (Fig. 5c). In the absence of depletion by antibodies^35, 43^, virus can persist in lung tissue for several days even in the absence of infection, and thus contribute to plaque assay titers, any significant decrement is encouraging as a potential treatment. More importantly in this model is the histological examination of the lungs for large hemorrhages related to actual SARS-CoV-2 infection, reflecting direct blood vessel damage as also seen in human lungs with fatal COVID-19^44^. In this regard, on histological examination, NMT5 virtually eliminated large COVID-19-related hemorrhages in the lungs of infected hamsters compared to vehicle when examined up to 5 days after infection (Fig. 5d). This translational model revealed a striking absence of large SARS-CoV-2-induced hemorrhages in the lungs of NMT5-treated hamsters vs. controls, with all controls displaying such hemorrhages while no NMT-5 treated animals did so (*n* = 12, *P* < 0.01 by Fisher Exact Test)^45^. While some inflammatory changes were noted in the NMT5-treated lung tissue compared to uninfected controls, it was far less than in the infected/untreated tissue (Fig. 5d), and the lethal effect of large hemorrhagic conversion was completely prevented by NMT5.

These findings benchmark favorably against our group’s published studies on the same model using antibodies directed against Spike protein^35, 43^. Additionally, proinflammatory cytokine and chemokine activation downstream of infection were abated with NMT5 treatment compared to vehicle control, as demonstrated by immunohistochemical staining of lung sections (Fig. 5e).

## Discussion

In summary, development of an oral drug to combat acute SARS-Cov-2 infection remains a high priority to treat the COVID-19 pandemic, particularly for the unvaccinated segment of the world population. Our findings provide proof that the cellular receptor of SARS-CoV-2, ACE2, can be S-nitrosylated to inhibit binding of SARS-CoV-2 Spike protein, thus inhibiting viral entry, infectivity, and cytotoxicity. Taking advantage of this finding, we developed a novel aminoadamantane nitrate compound, NMT5, that provides inhibition of SARS-CoV-2 activity by protein S-nitrosylation with a nitro group that is targeted to ACE2 by aminoadamantane-mediated viroporin channel blockade of the E protein^2, 3, 10–12^. The discovery that ACE2 could be S-nitrosylated was quite unexpected, as most authorities had postulated that the beneficial effects of NO on COVID-19 patients was due to a direct effect on the virus itself. These mechanistic insights should facilitate development of aminoadamantane nitrate drugs for acute antiviral therapy for human COVID-19. A key concept of this novel approach to ameliorating infection by SARS-CoV-2 is that these nitro-aminoadamantane compounds prevent the viral Spike protein from binding to the ACE2 receptor by S-nitrosylating the receptor in targeted fashion, facilitated by blockade of the vicinal viroporin E protein (Figs. 4, 5e, Extended Data Fig. 4). Hence, as our data indicate, drugs like NMT5 should also prevent new variants of the Spike protein from binding to ACE2 because ACE2 itself is blocked. In this manner, the aminoadamantane nitrate approach to COVID-19 drug therapy complements vaccine and antibody therapies, which are dependent on Spike protein antigenic sites and thus may eventually be susceptible to evasion by further Spike protein mutation. Critically, the binding of NMT5 to the viroporin channel also confers the ability to block spread of SARS-CoV-2 from one host to another. Mechanistically, NMT5 binds to the E protein viroporin channel on SARS-CoV-2 and then transfers NO to ACE2 on the host cell to prevent infection (**Fig. 4a, b**).

However, if a patient is already infected and takes NMT5, the newly produced viral particles will bind NMT5 via their E protein viroporin channels and hence viral infectivity will be limited when a new host is exposed to this virus because the new host’s ACE2 target protein will be S-nitrosylated by the drug attached to the viral particles as the virus approaches ACE2 on the new host.

## Online content

Any methods, additional references, Nature Research reporting summaries, source data, extended data, supplementary information, acknowledgements, peer review information; details of author contributions and competing interests; and statements of data and code availability are available at https://doi.org/xxxxx/xxxxxxx

## Methods

### Cell lines

HEK293T (System Biosciences, LV900A-1) and HEK293-Spike cells (SARS-CoV-2 Spike (D614)-expressing 293 cells [293-SARS2-S cells, InvivoGen]) were maintained in Dulbecco’s modified Eagle’s medium (DMEM) with GlutaMAX™ (DMEM, high glucose, GlutaMAX™ Supplement, Life Technologies, 10566016) supplemented with 10% fetal bovine serum (FBS; Sigma, F7524), 100 IU/ml, and 100 µg/ml penicillin-streptomycin (Thermo Fisher Scientific, 10378016) at 37 °C in a 5% CO_2_ incubator. Transfections were carried out with Lipofectamine 2000 (Life Technologies, 11668019) using the protocol recommended by the manufacturer. HeLa-ACE2 cells were a gift from David Nemazee (Scripps Research)^35^. Monkey Vero E6 cells (ATCC CRL-1586) were maintained in complete DMEM (Corning, 15-013-CV) containing 10% FBS, 1×PenStrep (Corning 20-002-CL), 2 mM L-glutamine (Corning, 25-005-CL) at 37 °C in a 5% CO_2_ incubator.

### Plasmids

hACE2 was a gift from Hyeryun Choe (Addgene plasmid #1786; http://n2t.net/addgene:1786 ; RRID:Addgene_1786)^46^. The C262A, C498A, C261/498A mutant ACE2 constructs were generated using the QuikChange Lightning Multi Site-Directed Mutagenesis Kit (Agilent Technologies, 210514) according to the manufacturer’s protocol. pGBW-m4252984 (SARS-CoV-2 E [envelope]) was a gift from Ginkgo Bioworks (Addgene plasmid #153898; http://n2t.net/addgene:153898; RRID:Addgene_153898). MLV-gag/pol, MLV-CMV-Luciferase, SARS-CoV-2, and VSV-G plasmids were a gift from David Nemazee (Scripps Research)^35^.

### Aminoadamantane and aminoadamantane nitrate drugs

Aminoadamantane nitrate compounds (blindly coded NMT2, NMT3, NMT5-NMT9, and NMT5-Met (metabolite, sans nitro group) were synthesized by and obtained from EuMentis Therapeutics, Inc. (Newton, MA), and have been described previously^6–9, 33^. The aminoadamantane compounds memantine and amantadine (blindly coded NMT1 and NMT4) were also obtained from EuMentis Therapeutics, Inc. All compounds were sent to Scripps Research for testing in a masked fashion, and compound identities were not revealed until after experiments were completed and analyzed blindly.

### Biotin-switch assays and immunoblotting

For analysis of S-nitrosylated proteins, we performed the biotin-switch assay as previously described ^47–49^. In brief, cells or tissue samples were lysed with HENTS buffer (100 mM Hepes, pH 7.4, 1 mM EDTA, 0.1 mM neocuproine, 0.1% SDS, and 1% Triton X-100) containing 10 mM methyl methanethiosulfonate [MMTS]). SDS solution (2 % w/v) was added to lysed samples to a final concentration of 1% and incubated for 20 minutes at 45°C with frequent vortexing to facilitate blockade of free thiol groups. After removing excess MMTS by acetone precipitation, S-nitrosothiols were reduced to thiols with 20 mM ascorbate. Newly formed thiols were then linked with the sulfhydryl-specific biotinylating reagent N-[6-biotinamido]-hexyl]-l′-(2′pyridyldithio) propionamide (Biotin-HPDP; Dojindo, SB17-10). Unreacted biotin-HPDP was removed by acetone precipitation, and the pellet was resuspended in HENS buffer (100 mM Hepes, pH 7.4, 1 mM EDTA, 0.1 mM neocuproine, 1% SDS), neutralized, and centrifuged to clear any undissolved debris. Five percent of the supernatant was used as the input for the loading control. Biotinylated proteins were pulled down with High Capacity NeutrAvidin-Agarose beads (Thermo Scientific, 29202) and analyzed by immunoblotting for S-nitrosylated ACE2, TMPRSS2, or Spike (S) protein. Protein samples were subjected to Bolt Bis-Tris Plus (Thermo Fisher Scientific, NW00102BOX) gel electrophoresis and transferred to PVDF membranes (Millipore, IPFL00010). Membranes were blocked with Odyssey blocking buffer (Li-Cor, 927-40000) for 30 minutes at RT and then probed with primary antibodies against ACE2 (1:3000, Abcam, ab15348; 1:3000, Proteintech, 21115-1-AP), TMPRSS2 (1:1000, Santa Cruz, sc-515727), or SARS-CoV-2 Spike protein (1:2000, Abcam, ab275759). After incubation with secondary antibodies (IR-dye 680LT-conjugated goat anti-mouse [1:20,000; Li-Cor, 926-68020] or IR-dye 800CW-conjugated goat anti-rabbit [1:15,000; Li-Cor, 926-32211]), membranes were scanned with an Odyssey infrared imaging system (Li-Cor). Image Studio (Li-Cor) software was used for densitometric analysis of immunoblots.

### Immunocytochemistry for SARS-CoV-2 Spike protein

Purified recombinant SARS-CoV-2 Spike (S1+S2) protein (10 μg/ml, Sino Biological, 40589-V08B1) exposed cells were fixed with 4% PFA for 15 minutes at RT, washed 3 times with PBS, and blocked (3% BSA, 0.3% Triton X-100 in PBS) for 30 minutes at RT. Cells were incubated with anti-SARS-CoV-2 Spike protein antibodies (1:200, Sino Biological, 40150-R007) overnight at 4 °C, followed by incubation with Alexa Fluor 488-conjugated secondary antibody. Cells were counterstained with 1 µg/ml Hoechst dye 33342 (Invitrogen). Cell images were acquired with a Nikon A1 Confocal Microscope using a 20 x/0.75 air objective (1 μm Z-stack). Maximum intensity projection of images was generated, and fluorescence intensity was analyzed with ImageJ software (https://imagej.nih.gov/ij/download.html) as previously described^50^.

### Expression and purification of human ACE2 protein

The N-terminal peptidase domain of human ACE2 (residues 19 to 615, GenBank: BAB40370.1) was cloned into phCMV3 vector and fused with C-terminal His-tag. The plasmid was transiently transfected into Expi293F cells using ExpiFectamine™ 293 Reagent (Thermo Fisher Scientific) according to the manufacturer’s instructions. The supernatant was collected at day 7 post-transfection. The His-tagged ACE2 protein was then purified by Ni-NTA (QIAGEN) affinity purification, followed by size exclusion chromatography. The ACE2 preparation then was buffer-exchanged to 1x PBS for the S-nitrosylation assay.

### Co-immunoprecipitation experiments

Cultured cells were harvested and lysed in 1% Triton X-100 in PBS. Equivalent protein quantities were immunoprecipitated with anti-ACE2 antibody (Abcam, ab15348)-conjugated magnetic beads (Dynabeads™ Protein A; Thermo Fisher Scientific, 10002D) for 90 minutes at RT. Immunoprecipitants were eluted and subjected to immunoblotting with anti-ACE2 antibody (1:1000, Cell Signaling, #15983) and anti-SARS-CoV-2 Spike protein antibody (1:2000, Abcam, ab275759).

### Mass spectrometry analysis of S-nitrosylated ACE2 protein

Biotin switch was performed as described above. Biotinylated proteins were then precipitated with iced acetone, pelleted, and solubilized in HENS buffer (100 mM Hepes, pH 7.4, 1 mM EDTA, 0.1 mM neocuproine, 1% SDS). The samples were desalted using a ZebaSpin desalting column (Thermo Scientific) pre-equilibrated with PBS, and biotinylated ACE2 protein was immunoprecipitated as described above. Immunoprecipitated ACE2 was eluted in 1% SDS solution and precipitated using methanol-chloroform. Dried pellets were dissolved in 8 M urea/100 mM triethylammonium bicarbonate (TEAB, pH 8.5). Samples were diluted to 2 M urea/100 mM TEAB, and proteins were trypsin digested overnight at 37 °C. The digested ACE2 peptides were enriched a second time by biotin-avidin affinity to enrich biotinylated peptides representing the initial SNO sites. After avidin enrichment, peptides were eluted by reduction using tris(2-carboxyethyl)phosphine (TCEP).

Samples were analyzed on a Thermo Orbitrap Eclipse mass spectrometer (Thermo). Samples were injected directly onto a 25 cm, 100 μm ID column packed with ethylene bridged hybrid (BEH) 1.7 μm C18 resin (Waters). Samples were separated at a flow rate of 300 nl/min on a nLC 1200 (Thermo) using a gradient of solution A (0.1% formic acid in water, 5% acetonitrile) and solution B (80% acetonitrile/0.1% formic acid). Specifically, a gradient of 1–25% B over 100 min, an increase to 40% B over 20 min, an increase to 100% B over another 10 min and held at 90% B for a 10 min was used for a 140 min total run time. Peptides were eluted directly from the tip of the column and nanosprayed directly into the mass spectrometer by application of 2.5 kV voltage at the back of the column. The Orbitrap Eclipse mass spectrometer was operated in data dependent mode. Full MS1 scans were collected in the Orbitrap at 120k resolution. The cycle time was set to 3 s, and within this 3 s, the most abundant ions per scan were selected for high energy collisional dissociation (HCD) with detection in the Orbitrap. Monoisotopic precursor selection was enabled and dynamic exclusion was used with exclusion duration of 5 s.

Protein and peptide identification were done with Integrated Proteomics Pipeline – IP2 (Integrated Proteomics Applications). Tandem mass spectra were extracted from raw files using RawConverter^51^ and searched with ProLuCID^52^ against Uniprot human database. The search space included all fully-tryptic and half-tryptic peptide candidates. Data were searched with 50 ppm precursor ion tolerance and 600 ppm fragment ion tolerance. Identified proteins were filtered to 10 ppm precursor ion tolerance using DTASelect ^53^ utilizing a target-decoy database search strategy to control the false discovery rate to 1% at the protein level^54^.

### Molecular dynamics simulations

The fully glycosylated, Cys^261^/Cys^498^-S-nitrosylated model of human ACE2 dimer bound to two SARS-CoV-2 Spike’s receptor binding domains (RBD) (with a 1:1 stoichiometry) is based on the cryo-EM structure of the ACE2/RBD/B0AT1 complex (PDB ID: 6M17)^27^. B0AT1 dimer chaperone coordinates were manually removed and N-glycans were added on both ACE2 and RBD in the same fashion as in Barros et al.^26^. The side chain of Cys^261^ and Cys^498^ was S-nitrosylated in both ACE2 protomers using Schrödinger Maestro (Schrödinger Release 2020-4: Maestro, Schrödinger, LLC, New York, NY, 2020.). ParamChem web interface was used to generate CHARMM36 suitable parameters for the S-N=O moiety^5^^5–58^. The glycosylated and S-nitrosylated ACE2/RBD construct was inserted into a lipid bilayer patch of 350 Å x 350 Å with a composition similar to that of mammalian cell membranes (56% POPC, 20% CHL, 11% POPI, 9% POPE, and 4% PSM)^59, 60^. Finally, the resulting system was embedded into an orthorhombic box of explicit waters and Na^+^/Cl^-^ ions at a concentration of 150 mM. Molecular dynamics (MD) simulations were performed on the Frontera supercomputer at the Texas Advanced Supercomputing Center (TACC) using NAMD 2.14^61^ and CHARMM36m all-atom additive force fields^62–64^. Excluding initial minimization and equilibration, a total of 310 ns were collected for analysis. We note that, except for Cys^261^ and Cys^498^ S-nitrosylation, the model described here is the same as the ACE2/RBD complex presented in Barros et al.^26^. Therefore, we refer to that work for a complete description of system setup procedures and simulation protocol.

Analysis of center of mass (COM) distance was performed with compute_center_of_mass function within MDtraj^65^. COM for each ACE2 protomer was calculated taking into account the amino acid backbones of residues 18-742. The distance between COMs was evaluated at each frame along a 310 ns trajectory for both the WT ACE2/RBD complex by Barros et al.^26^ and the model of the S-nitrosylated-ACE2/RBD complex presented here. As a reference, the distance between COMs from the cryo-EM structure (PDB: 6M17) was also calculated. The simulations were visually inspected using VMD^66^.

### SARS-CoV-2 virus generation

Monkey Vero E6 cells were plated in a T225 flask with complete DMEM containing 10% FBS, 1×PenStrep, 2 mM L-glutamine and incubated for overnight at 37 °C in a humified atmosphere of 5% CO_2_. The medium in the flask was removed, and 2 ml of complete DMEM containing the WA1 strain of SARS-CoV-2 (USA-WA1/2020 [BEI Resources, NR-52281)) were added to the flask at a multiplicity of infection (MOI) of 0.5. After incubation for 30 minutes at 34 °C in a 5% CO_2_ incubator, 30 ml of complete DMEM were added to the flask. The flask was then placed in a 34 °C incubator with 5% CO_2_ for 5 days. On day-5 postinfection, the supernatant was harvested and centrifuged at 1,000×g for 5 minutes. The supernatant was filtered through a 0.22 µM filter and stored at -80 °C.

### SARS-CoV-2/HeLa-ACE2 high-content imaging assay for infection

Control compounds solvated in DMSO were transferred into 384-well µclear-bottom plates (Greiner, Part. No. 781090-2B) using the ADS Labcyte Echo liquid handler. Aminoadamantane and aminoadamantane nitrate compounds to be screened were solvated in saline solution on ice immediately before use and transferred into the assay plates in 5 µl DMEM with 2% FBS (assay medium). HeLa-ACE2 cells were added to plates in assay medium at a density of 1.0×10^3^ cells per well to a 13 µl volume. Plated cells were transported to the BSL3 facility at Scripps Research, and within 1 h, 13 µl SARS-CoV-2 diluted in assay media was added at a multiplicity of infection (MOI) = 0.65. Cells were incubated for 24 h at 34 ℃ in a 5% CO_2_ incubator and then fixed with 8% formaldehyde for 1 h. Human polyclonal sera diluted at 1:500 in Perm/Wash buffer (BD Biosciences, 554723) were added to the plate and incubated at room temperature (RT) for 2 h. Six µg/ml of goat anti-human H+L conjugated Alexa 488 (Thermo Fisher Scientific, A11013) together with 8 µM of antifade-46-diamidino-2-phenylindole (DAPI; Thermo Fisher Scientific D1306) in SuperBlock T20 (PBS) buffer (Thermo Fisher Scientific, 37515) were added to the plate and incubated at RT for 1 h in the dark. Four fields were imaged per well using the ImageXpress Micro Confocal High-Content Imaging System (Molecular Devices) with a 10× objective. Images were analyzed using the Multi-Wavelength Cell Scoring Application Module (MetaXpress) where DAPI staining was used to identify host-cell nuclei (the total number of cells in the images) and the SARS-CoV-2 immunofluorescence signal for identification of infected cells.

### Uninfected host cell cytotoxicity counterscreen

Compounds were prepared and plated in 384-well plates as for the infection assay. HeLa-ACE2 cells were seeded in the assay-ready plates at 1.6×10^3^ cells/well in assay medium, and plates were incubated for 24 h at 37 ℃ with 5% CO_2_. To assess cell viability, the Image-iT DEAD green reagent (Thermo Fisher, I10291) was used according to manufacturer’s instructions. Cells were fixed with 4% paraformaldehyde (PFA) and counterstained with DAPI. Fixed cells were imaged using the ImageXpress Micro Confocal High-Content Imaging System (Molecular Devices) with a 10× objective, and total live cells per well quantified in the acquired images using the Live Dead Application Module (MetaXpress).

### Data analysis of the compound screening results

The *in vitro* infection assay and the host cell cytotoxicity counterscreen data were uploaded to Genedata Screener, Version 16.0. Data were normalized to neutral (DMSO) minus inhibitor controls (2.5 µM remdesivir for antiviral effect, and 10 µM puromycin for infected host cell toxicity). For the uninfected host cell cytotoxicity counterscreen, 10 µM puromycin (Sigma) was used as a positive control. For dose-response experiments, compounds were tested in technical triplicates, and dose curves were fitted with the four parameter Hill Equation. Replicate data were analyzed using median condensing. The full dataset is supplied in Extended Data Table 1.

### Pseudoviral entry assay

To measure SARS-CoV-2 viral infectivity, we performed pseudoviral entry assays as previously described^35^. In brief, HEK293T cells were transiently co-transfected with MLV-gag/pol, MLV-CMV-Luciferase plasmid, and SARS-CoV-2 Spike (D614) or VSV-G plasmid. Two days later, supernatants containing pseudotyped virus particles were collected. To assay for pseudoviral entry, HeLa-ACE2 cells were seeded in 96-well plates at 10,000 cells/well (PerkinElmer Inc. 6005680). One day later, cells were incubated with diluted pseudovirus. After 48 h to allow for viral transduction, cells were lysed and assayed for luciferase activity by Steady-Glo^®^ (Promega, E2510) according to the manufacturer’s instructions. Luminescence was quantified using a Luminoskan Ascent plate leader (Thermo Fisher Scientific). To assess the ability of NMT5 to reduce viral infection in human-ACE2 cells with SARS-CoV-2 variants. Lenti-SARS-CoV-2 N501Y Spike, SARS-CoV-2 K417N, E484K, N501Y Spike, and VSV-G pseudoviral particles were obtained from the Rhode Island Hospital lentivirus construct core^67^, or Lenti-SARS-CoV-2 delta and omicron variant pseudoviral particles were obtained from the PBS Bioscience (#78215 and #78348, respectively). SARS-CoV-2 variant preparations were diluted in assay media and added at 4.0×10^6^ TU/1×10^4^ or 1.5×10^2^ TU/1×10^4^ cells (for delta variant), or 9×10^2^ TU/1×10^4^ cells (for omicron variant) in the presence or absence of 20 µM NMT5. Plates were centrifuged at 600×g for 1 h at RT, and then cells were incubated at 37 ℃ in a 5% CO_2_ incubator. After 48 h to allow for viral transduction, cells were lysed and assayed for luciferase activity by Steady-Glo®. Luminescence was quantified using a Luminoskan Ascent plate reader.

### Pharmacokinetic testing and analysis

Golden Syrian hamsters (110-150 gm, *n* =3 for each compound) were dosed by oral gavage at 10 mg/kg. Blood samples were obtained at 30 min, 1, 3, 7, 24, 32, and 48 h. The blood samples were collected in BD Vacutainers (BD, #366664) containing sodium heparin, and centrifuged. The processed plasma samples are stored at -20 °C until high-performance liquid performance-tandem mass spectrometry (HPLC-MS/MS) analysis. Animals were then sacrificed, and lung and kidney tissues harvested for biotin-switch analysis for SNO-ACE2. To quantify the test compound in the collected plasma samples, a plasma calibration curve was generated by spiking aliquots of drug-free plasma with the test compound at the specified concentration levels. Spiked and collected plasma samples were treated with an aliquot of acetonitrile containing a known concentration of an Internal Standard (IS). The extraction solvent was analyzed on an Agilent 1100 LC mated to a AB Sciex 4000 Q TRAP MS. Separation of the analytes was achieved by using a Phenomenex Kinetex EVO C18 50 x 2.1mm column (Phenomenex, 00B-4633-AN), and a mobile phase consisting of (A) 0.1% formic acid in water and (B) 0.1% formic acid in acetonitrile. The LC gradient consisted of a 0.5 ml/min flow rate starting at 100% of (A). The LC gradient then ramped to 90% (B) over 0.1 min, and held for 2.5 min. The gradient reverted to 100% (A) over 0.1 min and allowed for 2 min of re-equilibration time. Ionization spray (IS) voltage was set to 4000, with the source temp set at 400 °C. Analytes were monitored by Multiple Reaction Monitoring (MRM) in positive ionization mode. Peak areas were recorded, and the concentrations of the test compound in the unknown plasma samples were quantified by the calibration curve using Sciex Analyst software (PE Sciex). Phoenix WinNonLin 8.1 software (Certara) was used in NCA mode to determine the PK parameters from the LC-MS/MS measured values. NMT5 PK analysis was performed by “Serial Sampling” where we followed the PK of the drug in each individual animal (animals 71, 73, and 75), and then averaged those PK values. We took this approach because all the animals had values for the full-time course available, as detailed in Extended Data Table 3. For NMT3 PK, the “sparse sampling” procedure was utilized to account for plasma concentrations from animals that do not have a full-time course profile available (animals 372-374), in part due to the short half-life and rapid metabolism to the hydroxylated form with release of the nitrate group, as detailed in Extended Data Table 4. The difference in methodology results in “Serial Sampling” averaging the PK values and “Sparse Sampling” averaging the concentrations used to determine the PK values. Sparse sampling was also necessary because tissue samples were obtained at two of the timepoints (1 h and 24 h) so the study parameters had to be modified. Additionally, both parent drug (NMT3) and metabolite were detected, making the samples sparser. Summary PK data are shown in Extended Data Table 2, and the full datasets are supplied in Extended Data Table 3 (for NMT5) and Extended Data Table 4 (for NMT3).

### Syrian hamster COVID-19 model

Eight-week old Golden Syrian hamsters (110-150 gm, Charles River) challenged with a dose of 1 × 10^5^ or 10^6^ plaque forming units (PFU) of SARS-CoV-2 (USA-WA1/2020) by intranasal administration in a volume of 100 µl DMEM as previously described^35^. For the experiments shown, the hamsters were subsequently administered via oral gavage either vehicle or the aminoadamantane nitrate candidate drugs NMT3 or NMT5 in a volume of 500 µl, with the initial dose timed for delivery right after challenge with SARS-CoV-2 and a second dose administered 12 h later. At 2-d and 5-d postinfection, lung tissue was collected for assessing viral titers and histology as described^35^.

### Plaque assay for SARS-CoV-2 viral titers

SARS-CoV2 titers were measured 2 days and 5 days after infection by homogenizing hamster lungs in DMEM 2% FBS using 100 µm cell strainers (Myriad 2825-8367). Homogenized lungs were titrated 1:10 over 6 steps and layered over Vero cells. After 1 h of incubation at 37 °C, a 1% methylcellulose in DMEM overlay was added, and the cells were incubated for 3 days at 37 °C. Cells were then fixed with 4% PFA and plaques were counted by crystal violet staining and expressed as plaque forming units (PFU). PFU after treatment with NMT3 was statistically equal to that of vehicle control.

### Lung histology and immunohistochemistry

Lung tissue from Golden Syrian Hamsters was stored in zinc formalin for ∼72 h. The tissue was processed for paraffin embedding, and sections cut at a thickness of 5 µm. The tissue was then stained with hematoxylin and eosin (H&E). The slides were scanned at 20X using an Aperio AT2 whole slide scanner. For immunohistochemistry, lung sections were fixed with 4% PFA for 15 min and washed 3 times with PBS. Sections were blocked with 3% BSA and 0.3% Triton X-100 in PBS for 30 min. Sections were incubated with primary antibody in blocking solution overnight at 4 °C and then washed with PBS. The appropriate Alexa Fluor (488, 555) conjugated secondary antibodies were used at 1:500, plus Hoechst 33342, Trihydrochloride, Trihydrate dye (1:1,000, ThermoFisher Scientific, catalog #H3570) to visualize nuclei for 1 h at RT. Primary antibodies and dilutions were as follows: Mouse anti-TNFα (5 µg/ml, Abcam, #ab1793) and rabbit anti-macrophage inflammatory protein 1α (MIP-1α)/CCL3+CCL3L1 (1:250, Abcam, #ab259372). Slides were imaged using a C2 confocal microscope (Nikon) in a masked fashion.

### Patch-clamp analysis of SARS-CoV-2 envelope (E) protein viroporin ion channel activity

Using the patch-clamp technique, we recorded whole-cell currents from untransfected HEK293T cells and cells transfected with SARS-CoV-2 envelope (E) protein. Cells were recorded at RT in Hepes-supplemented Hanks’ balanced salt external solution with the following composition (in mM): NaCl 138, KCl 5.3, KH_2_PO_4_ 0.4, Na_2_HPO_4_ 0.3, NaHCO_3_ 4.2, MgCl_2_ 0.5, MgSO_4_ 0.4, CaCl_2_ 2.0, D-glucose 5.6 and Hepes 10; pH 7.4, 300 mOsm. Patch pipettes were filled with an intracellular pipette solution composed of (in mM): 113 Kgluconate, 6 KCl, 4.6 MgCl_2_, 1.1 CaCl_2_, 10 Hepes, 10 EGTA, 4 Na_2_-ATP, 0.4 Na_2_-GTP, pH adjusted to 7.3 with KOH; osmolality 291 mOsm. Whole-cell currents were recorded using a computer-controlled patch-clamp amplifier (Multiclamp 700B, Molecular Devices). Traces were filtered at 2 kHz and sampled at 20 kHz. Drugs were dissolved in water and stored as 100 mM stocks at -20 °C. During experiments, drugs were dissolved in extracellular solution and tested at concentrations of 5 and 10 µM. HEK293T cells were plated on 12-mm diameter glass coverslips coated with a mixture of rat tail type I collagen and poly-D-lysine. For transient expression in HEK293T cells, we used a transfection reagent (Fugene® HD, Promega) to co-transfect plasmids containing cDNAs for SARS-CoV-2 E protein (pGBW-m4133502, Addgene) and green fluorescent protein (GFP) at a ratio of 1:0.1 (0.5:0.05 µg/well, respectively).

### SNO-ACE2 targeting assay via E protein viroporin channel

SARS-CoV-2 E plasmid was transiently transfected into HEK293-Spike cells with Lipofectamine 2000 (controls received Lipofectamine 2000 vehicle alone); one day later, cells were harvested by tapping the plate in pre-warmed PBS. These cells were then added onto pre-plated HeLa-ACE2 cells in the presence or absence of NMT5. After 30 min, all cells were collected and subjected to biotin switch-assay and immunoblotting with anti-ACE2 antibody to assess the levels of SNO-ACE2 and total input ACE2.

### Quantification and statistical analysis

A Power Analysis of our prior data was used to determine the number of replicates needed for statistical purposes. The number of replicates or experiments is indicated in the individual figure legends. Data are expressed as mean ± s.e.m. Differences between experimental groups were evaluated using an ANOVA followed by a post hoc Tukey’s or Fisher’s LSD test for multiple comparisons, or a Student’s t test for comparison of two groups, except for the statistical assessment of lung hemorrhages in the Syrian hamster COVID-19 model, which employed a Fisher exact test with 2 x 2 contingency table. A *P* value < 0.05 was considered significant. Statistical analyses were performed using GraphPad Prism software.

### Reporting summary

Further information on research design is available in the Nature Research Reporting Summary linked to this paper.

### Data availability

All data are available in the main text or the supplementary materials. All plasmids generated in this study are available from S.A.L. under a material transfer agreement with The Scripps Research Institute.

## Acknowledgements

We thank David Nemazee (Scripps Research) for providing HeLa-ACE2 cells and plasmids for pseudovirus. This work was supported in part by NIH grants RF1 AG057409, R01 AG056259, R01 DA048882, R35 AG071734 and DP1 DA041722 (to S.A.L.), R01 AG061845 (to T.N.), UM1 AI144462 (to D.R.B.), P41 GM103533 (to J.R.Y.), HOPE T32 Training Grant T32AI007384 (to L.N.C.), California Institute for Regenerative Medicine (CIRM) grant DISC2 COVID19-11811, COVID-19 awards from Fast Grants (to S.A.L.), and grants from the Bill & Melinda Gates Foundation #OPP1107194 (to Calibr) and INV-004923 (to I.A.W). The molecular dynamics simulations were supported by NIH R01 GM132826, NSF RAPID (MCB-2032054), an award from the RCSA Research Corp. and a UC San Diego Moore’s Cancer Center 2020 SARS-CoV-2 seed grant (to R.E.A.), and the Interfaces Graduate Training Program, NIH T32 EB009380 (to M.A.R). We thank the Texas Advanced Computing Center (TACC) Frontera team and acknowledge computer time made available through a Director’s Discretionary Allocation (made possible by NSF award OAC-1818253). The Lentivirus Construct Core of the COBRE Center for Stem Cells and Aging was supported by NIH grant P20 GM119943.

## Author contributions

S.A.L. conceived, designed, and supervised the execution of the entire project. C.O., T.N., and S.A.L. formulated the detailed research plans, interpreted experimental results, and wrote the first draft of the manuscript. C.O. performed a majority of molecular and biochemical experiments, and X.Z. carried out biochemical experiments with mutant ACE2. N.B., S.R.M., N.S., D.R.B., and T.F.R. performed *in vivo* experiments with the Syrian hamster model. D.T. and L.N.C. performed the immunohistochemistry on hamster lung tissue. J.P.-C., M.T., and S.G. performed electrophysiology experiments. M.A.B., A.J.R., A.K.W., V.C., A.K.G., and A.K.C. conducted compound screening and pharmacokinetic analyses. T.N., J.K.D., and J.R.Y. conducted mass spectrometry assays. M.A.R., F.L.K., L.C., and R.E.A. conducted molecular dynamics simulations. H.L. and I.A.W. produced recombinant ACE2 and helped write the structure-related aspects of the manuscript. C.O., T.N., I.A.W., R.E.A., D.R.B., J.R.Y., T.F.R., A.K.C., and S.A.L. reviewed and edited the manuscript.

## Competing interests

The authors declare that S.A.L. is an inventor on patents for the use of memantine and various aminoadamantane nitrate compounds for neurodegenerative and neurodevelopmental disorders. He is also an inventor on composition of matter patents and use patents for aminoadamantane nitrate compounds in treating COVID-19 and other viral diseases. Per Harvard University guidelines, S.A.L. participates in a royalty-sharing agreement with his former institution Boston Children’s Hospital/Harvard Medical School, which licensed the drug memantine (Namenda®) to Forest Laboratories, Inc./Actavis/Allergan/AbbVie for use in dementia. The aminoadamantane nitrate compounds have been licensed to EuMentis Therapeutics, Inc. (Newton, Massachusetts). C.B. is a chemist employed at EuMentis Therapeutics. The other authors declare no financial conflicts of interest relevant to this publication.

## Additional information

**Extended data** is available for this paper at https://doi.org/xxxxx/xxxxx.

**Supplementary information** The online version contains supplementary material available at https://doi.org/xxxxxx.

**Correspondence and requests for materials** should be addressed to Stuart A. Lipton.

**Peer review information** *Nature Microbiology* thanks the anonymous reviewers for their contribution to the peer review of this work.

**Reprints and permissions information** is available at http://www.nature.com/reprints.

---

**Extended Data Fig. 1.**
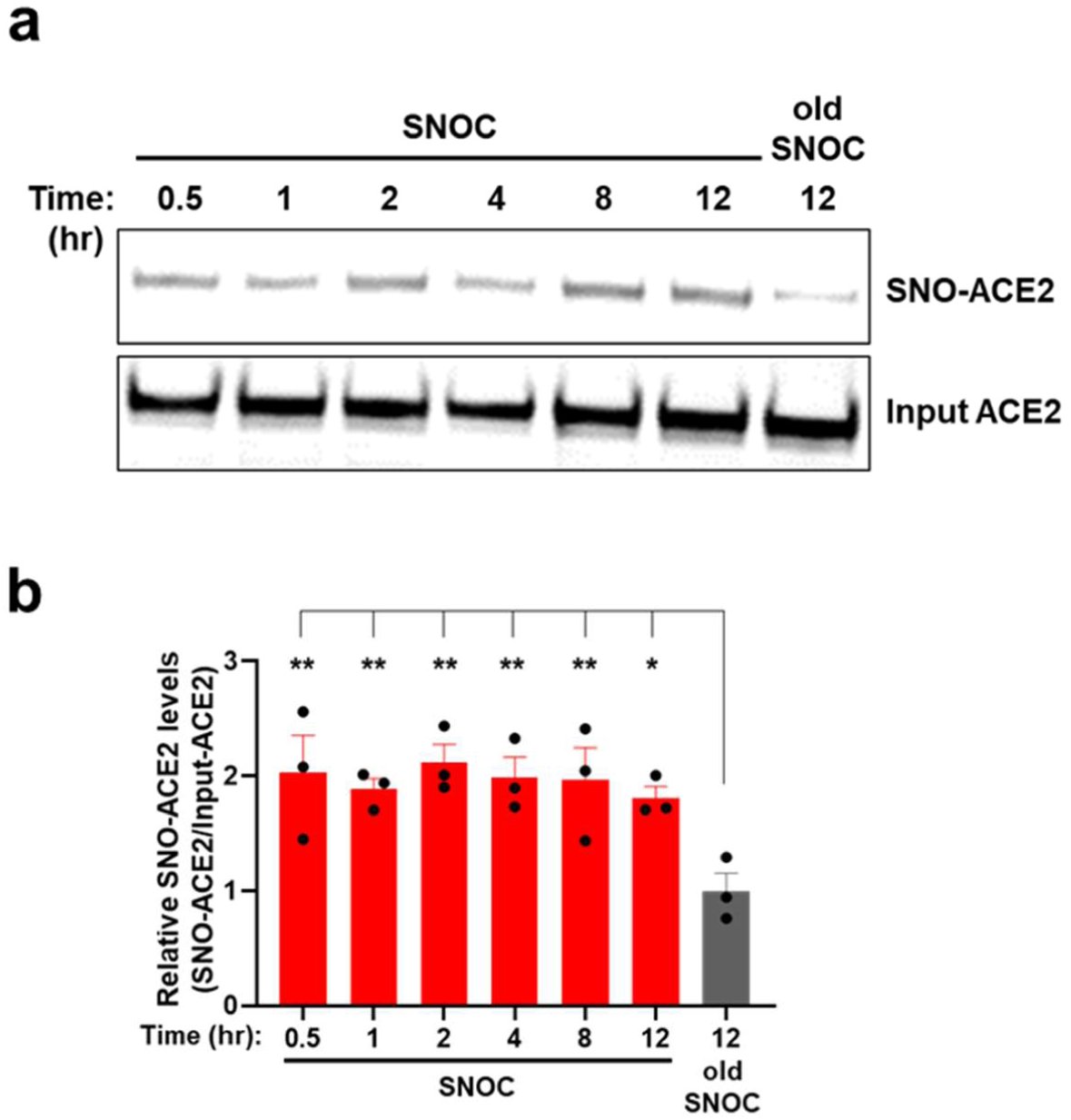
S-Nitrosylation of ACE2 persists for at least 12 hours. **a**, HELA-ACE2 cells were exposed to 100 μM SNOC; 30 min later the cells were incubated in serum free medium for the time periods indicated. Cell lysates were then subjected to biotin-switch assay to assess protein S-nitrosylation, which was detected by immunoblotting with anti-ACE2 antibody. **b**, Ratio of SNO-ACE2/input ACE2 protein. Data are mean + s.e.m., \**P* < 0.05 by ANOVA with Fisher’s LSD multiple comparisons. *n* = 3 biological replicates.

**Extended Data Fig. 2.**
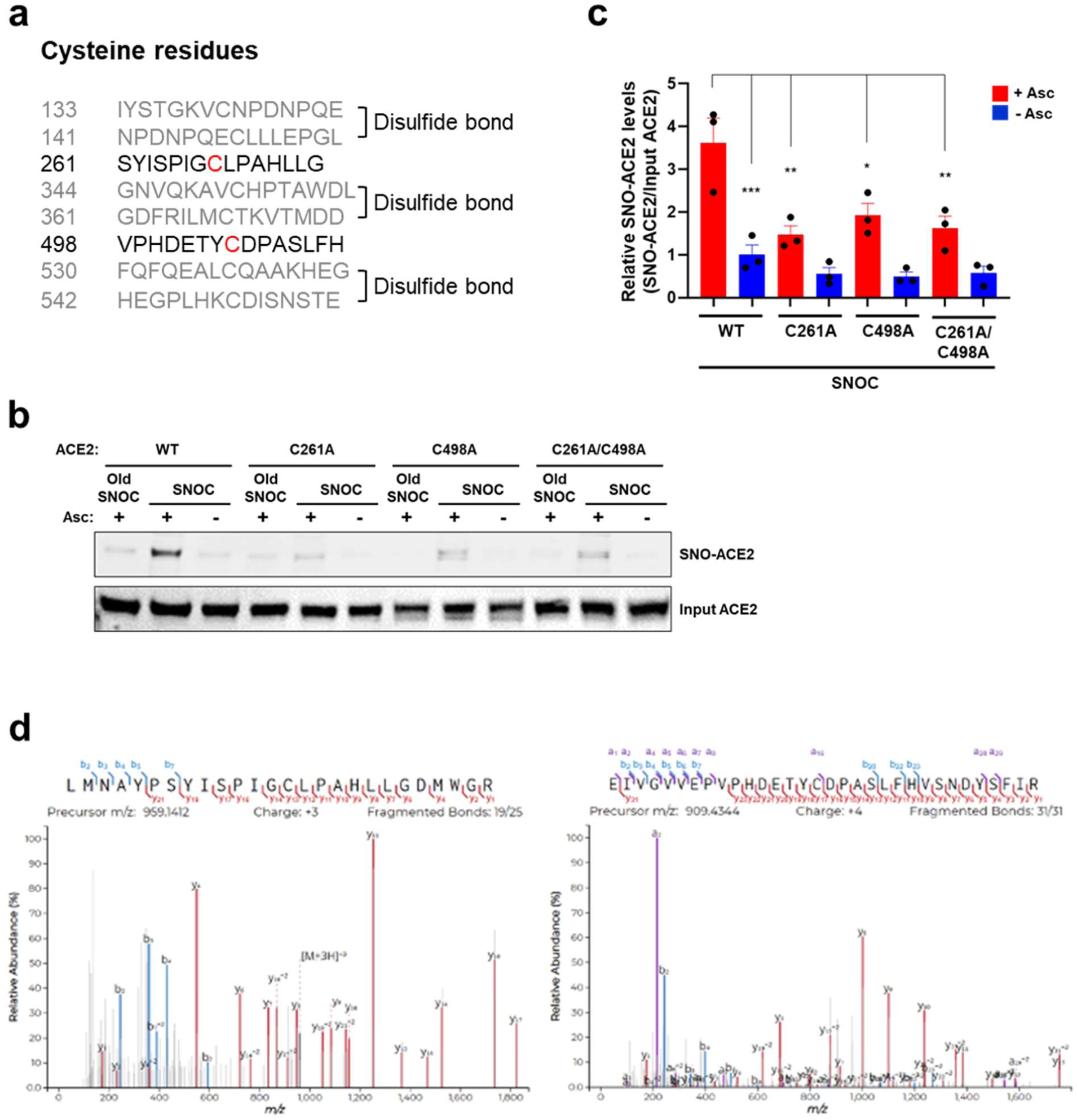
Identification of cysteine residues in ACE2 that are S-nitrosylated. **a**, List of human ACE2 peptides (± 7 amino acid residues flanking a cysteine residue); gray: peptides involved in disulfide bond formation; black: peptides containing a free cysteine thiol (red) that could potentially be S-nitrosylated. **b**, HEK293T cells were transiently transfected with plasmids containing human WT ACE2 or cysteine mutant ACE2 (C261A, C498A, or C261/498A). One day after transfection, cells were exposed to 100 μM SNOC. After 20 minutes, cells were subjected to biotin-switch assay. Absence of ascorbate (Asc -) served as a negative control. **c**, Ratio of SNO-ACE2/input ACE2. Data are mean + s.e.m., \**P* < 0.05, \*\**P* < 0.01, \*\*\**P* < 0.001 by ANOVA with Tukey’s multiple comparisons. *n* = 3 biological replicates. **d**, HEK293T cells expressing ACE2 were exposed to SNOC and subjected to biotin switch. The peptides were eluted by reduction for subsequent detection by LC-MS/MS. Representative MS/MS spectra of detected peptides from human ACE2 containing Cys^261^ (*left*) or Cys^498^ (*right*) are shown.

**Extended Data Fig. 3.**
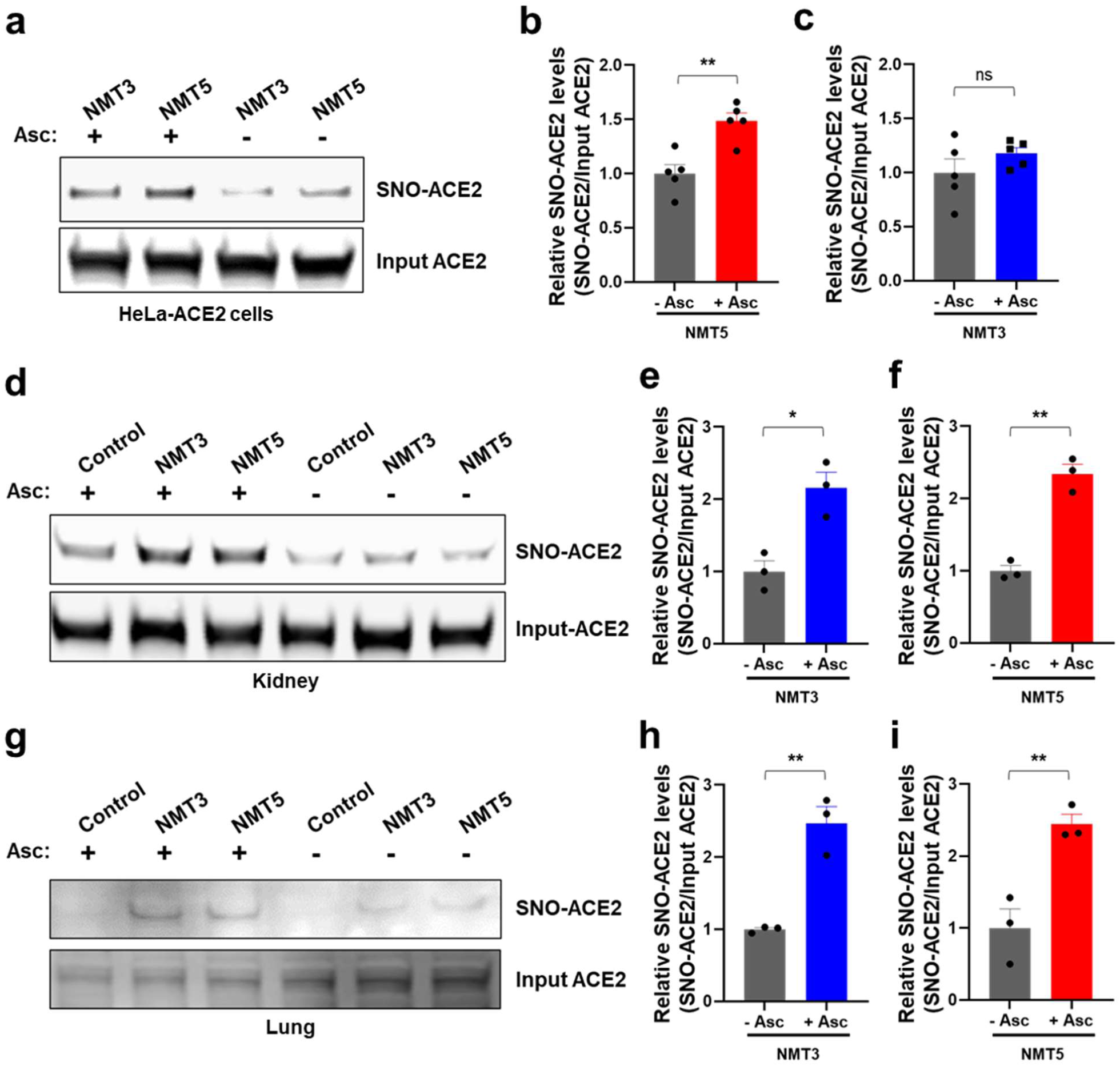
NMT5 S-nitrosylates ACE2 *in vitro* and *in vivo.* **a**, Detection of SNO-ACE2 *in vitro*. HeLa-ACE2 cells were exposed to 10 μM NMT3 or 5 μM NMT5. After 1 h, cells were subjected to the biotin-switch assay in the presence or absence of ascorbate. SNO-ACE2 and input ACE2 were detected by immunoblotting with anti-ACE2 antibody. **b**, **c**, Ratio of SNO-ACE2/input ACE2. Data are mean ± s.e.m., \*\**P* < 0.01 by two-tailed Student’s *t* test. ns: not significant, *n* = 5 biological replicates. **d**–**i**, Detection of SNO-ACE2 *in vivo*. Syrian hamsters received 10 mg/kg of NMT3 or of NMT5 by oral gavage and were sacrificed 48 h later. Kidney and lung tissues were subjected to biotin-switch assay in the presence or absence of ascorbate. Note that in some samples, low levels of SNO-ACE2 were observed in control tissue, suggesting endogenous S-nitrosylation of ACE2 may occur at low levels. Graphs show ratio of SNO-ACE2/input ACE2. Data are mean + s.e.m., \**P* < 0.05, ** *P* < 0.01 by two-tailed Student’s *t* test. *n* = 3 Syrian hamsters for each condition.

**Extended Data Fig. 4.**
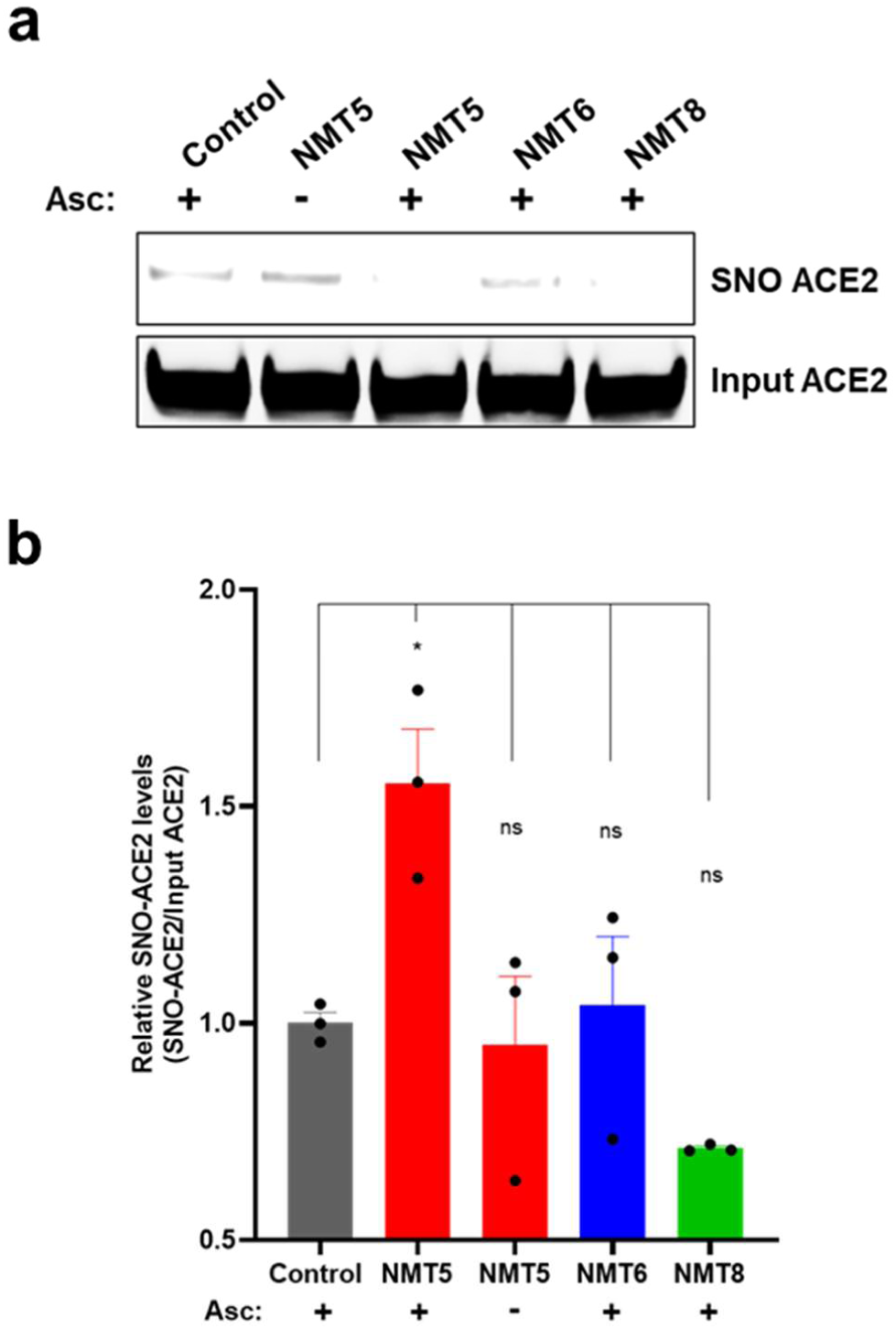
Protein S-nitrosylation of ACE2 by NMT5. **a**, HeLa-ACE2 cells were treated with 10 μM NMT5, NMT6, or NMT8. After 1 h, cell lysates were subjected to biotin-switch assay for protein S-nitrosylation, detected by immunoblotting with anti-ACE2 antibody. The ascorbate minus (Asc-) sample served as a negative control. **b**, Ratio of SNO-ACE2/input ACE2 protein. Data are mean + s.e.m., \**P* < 0.05, ns: not significant by ANOVA with Tukey’s multiple comparisons. *n* = 4 biological replicates.

**Extended Data Fig. 5.**
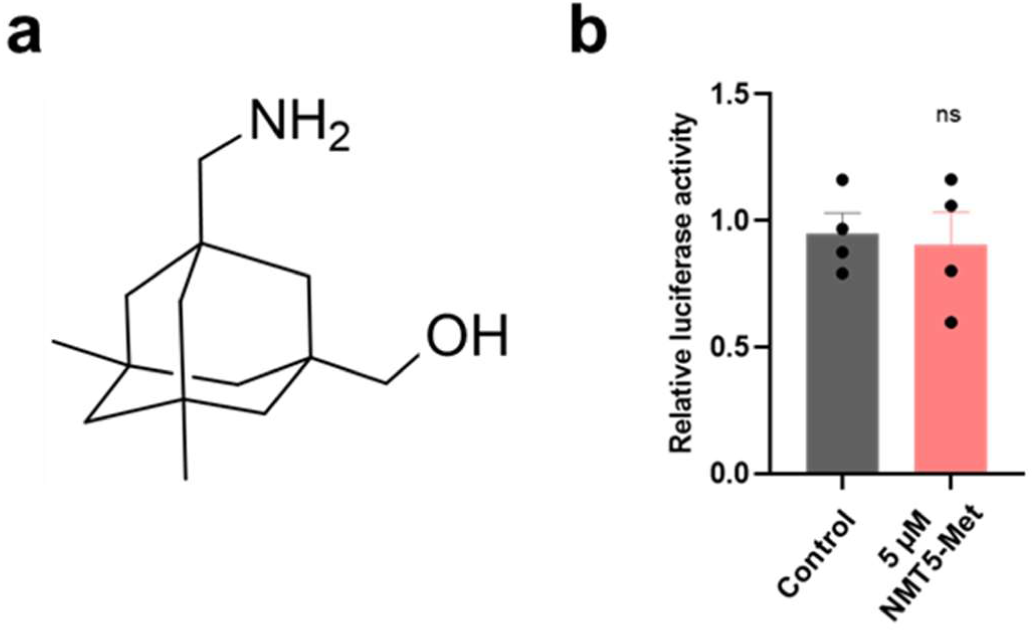
Critical role of nitro group of NMT5 suppressing SARS-CoV-2 infection on pseudovirus entry assay. **a**, Chemical structure of NMT5 metabolite (NMT5-Met, lacking the nitro group). **b**, HeLa-ACE2 cells were incubated in the presence and absence of 5 µM NMT5-Met with SARS-CoV-2 Spike (D614) pseudovirus particles. After 48 h, viral transduction efficiency was monitored by luciferase activity. Inhibitory activity was lost in the absence of the nitro group (compare with Fig. 3c, e). Data are mean + s.e.m., ns: not significant by two-tailed Student’s *t* test. *n* = 4 biological replicates.

**Extended Data Fig. 6.**
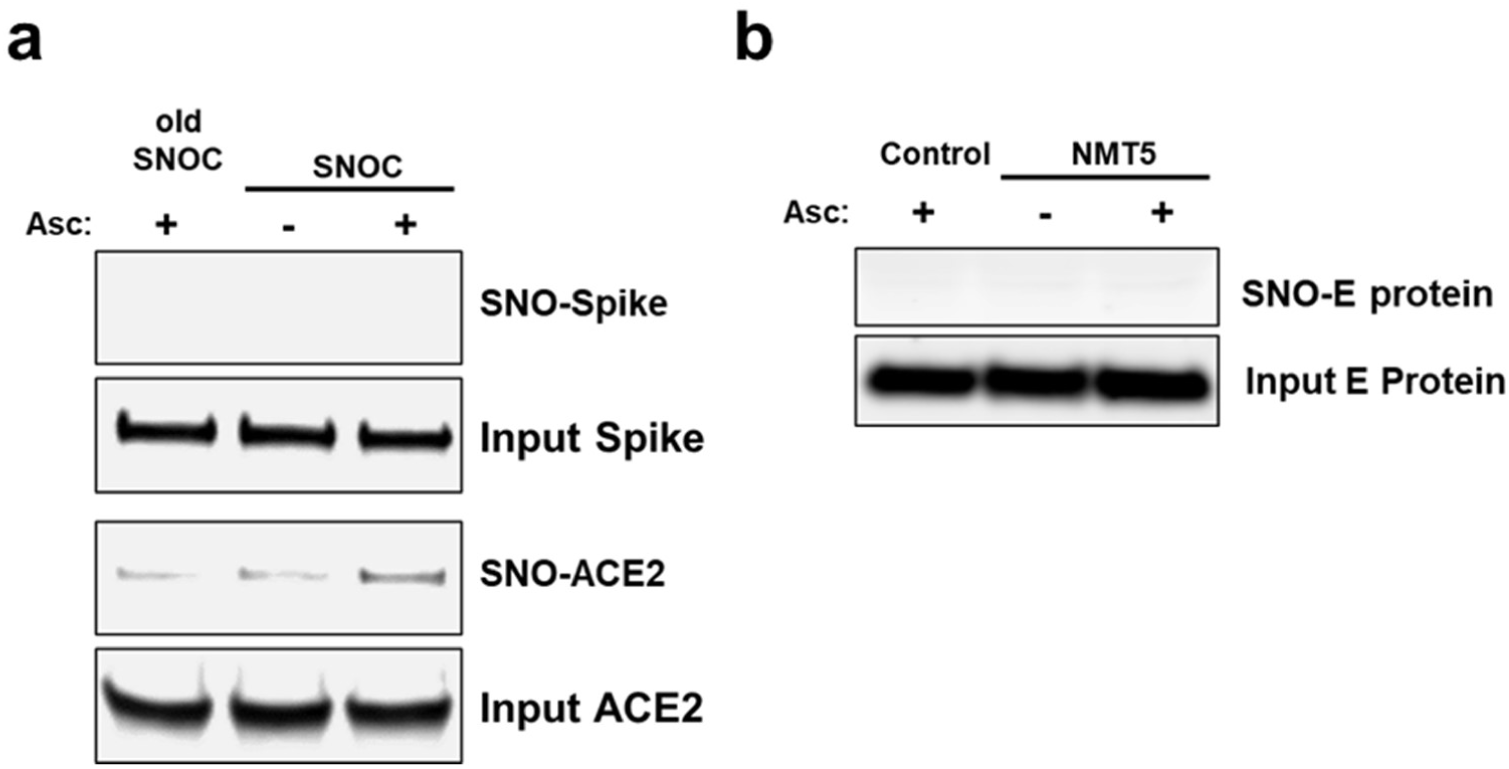
Lack of S-nitrosylation of Spike protein and E protein. **a**, Purified recombinant SARS-CoV-2 Spike (S1+S2) protein and ACE2 protein were exposed to 100 μM SNOC; 30 min later, samples were subjected to biotin-switch assay in the presence or absence of ascorbate (Asc) to assess protein S-nitrosylation. **b**, Lack of E protein S-nitrosylation by NMT5. HA-tagged E protein plasmid was transiently transfected into HEK293 cells. One day after transfection, cells were exposed to 10 μM NMT5. After 1 hour, the cells were harvested and subjected to biotin-switch assay in the presence or absence of ascorbate (Asc) to assess protein S-nitrosylation, which was detected by immunoblotting with anti-HA antibody.

**Extended Data Table 1.** Dose-response of drugs tested in SARS-CoV-2/HeLa-ACE2 high-content imaging and uninfected cytotoxicity assays. Dose-response analysis of *in vitro* infection assay and uninfected host cell cytotoxicity data for control (apilimod, remdesivir, puromycin) and test compounds, including dose-response curves and curve fit parameters. For the infection assay two assay metrics (% infected cells and total cells per well) are reported. This table is located in a separate EXCEL file.

**Extended Data Table 2.**
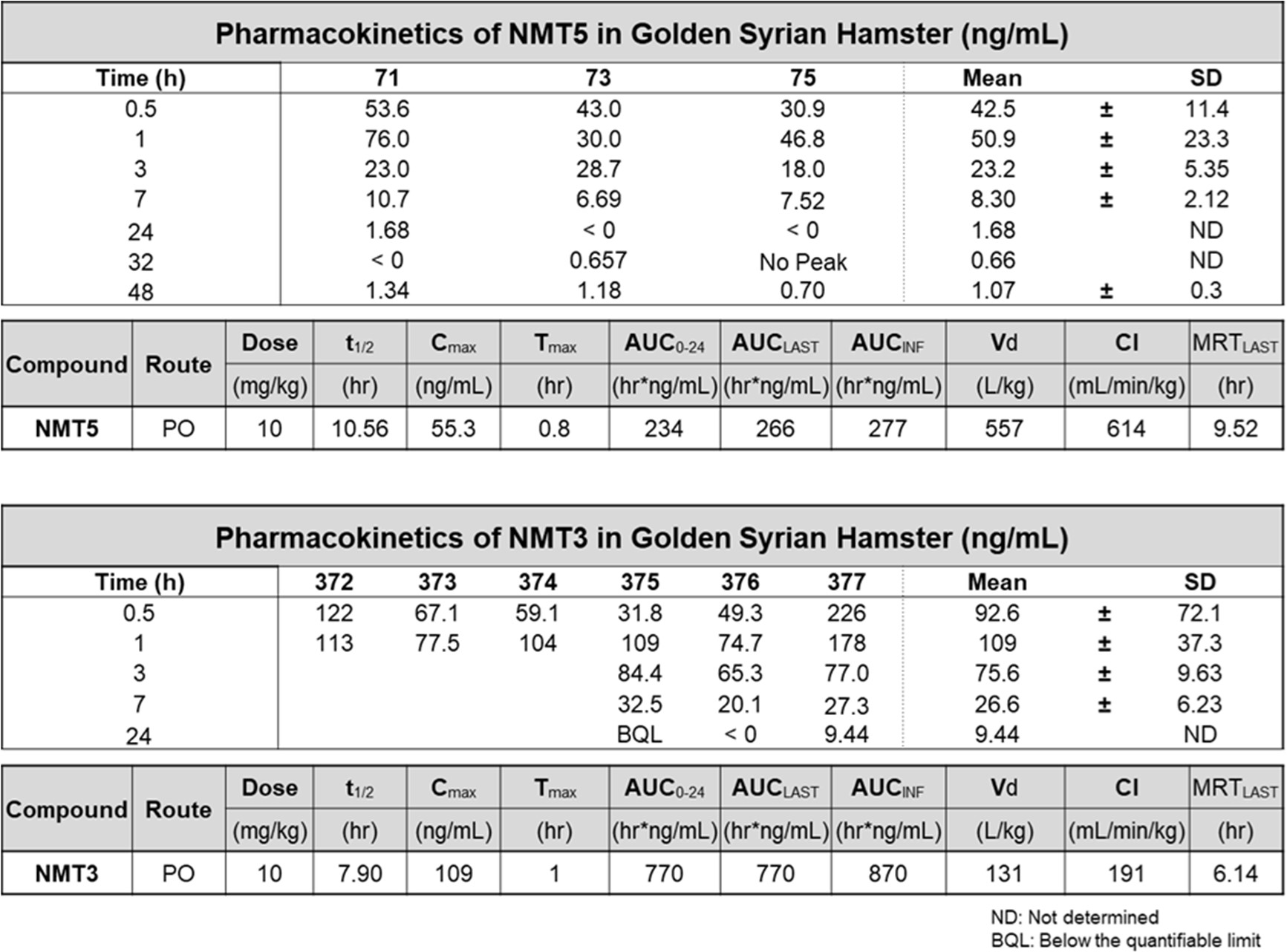
Summary values of PK data for NMT5 and NMT3. Plasma concentrations measured by LC-MS/MS of NMT5 or NMT3 in Syrian hamsters collected at timepoints ranging from 0.5 hours to 48 hours after a single 10 mg/kg dose administered by oral gavage. Pharmacokinetic (PK) analysis of NMT5 (*n* = 3) and NMT3 (*n* = 6) in hamsters was assessed using non-compartmental analysis (NCA) of extrapolated plasma concentrations over time using Phoenix WinNonLin software.

**Extended Data Table 3.** PK data for NMT5. Full dataset for plasma concentrations of NMT5 and metabolite quantified by LC-MS/MS after a single 10 mg/kg dose administered by oral gavage to Syrian hamsters. Extrapolation of plasma concentrations was determined from an NMT5-spiked standard curve with a linear range of 0.3 ng/ml - 1250 ng/ml, and a standard curve with a linear range of 0.3 ng/ml - 5000 ng/ml for the metabolite. Determination of PK parameters was conducted by NCA analysis of serially collected samples using Phoenix WinNonLin software. This table is located in a separate EXCEL file with multiple tabs accessible by clicking on the bottom of the page.

**Extended Data Table 4.** PK data for NMT3. Full dataset for plasma concentrations of NMT3 and metabolite quantified by LC-MS/MS after a single 10 mg/kg dose administered by oral gavage to Syrian hamsters. Extrapolation of plasma concentrations was determined from an NMT3-spiked standard curve with a linear range of 4.88 ng/ml – 10,000 ng/ml. The metabolite for NMT3 was quantified by extrapolation from a standard curve with a linear concentration range of 2.44 ng/ml – 10,000 ng/ml. Determination of limited PK parameters was conducted by NCA analysis of sparsely collected samples using Phoenix WinNonLin software. This table is located in a separate EXCEL file with multiple tabs accessible by clicking on the bottom of the page.

